# A Deep Learning approach predicts the impact of point mutations in intronic flanking regions on micro-exon splicing definition

**DOI:** 10.1101/868786

**Authors:** Lucas F. DaSilva, Ana C. Tahira, Vinicius Mesel, Sergio Verjovski-Almeida

## Abstract

While mammalian exons are on average 140-nt-long, thousands of human genes harbor micro-exons (≤ 39 nt). Large numbers of micro-exons have their splicing altered in diseases such as autism and cancer, and yet there is no systematic assessment of the impact of point mutations in intronic flanking-sequences on the splicing of a neighboring micro-exon. Here, we constructed a model using the Convolutional Neural Network (CNN) to predict the impact of point mutations in intronic-flanking-sequences on the splicing of a neighboring micro-exon. The prediction model was based on both the sequence contents and conservation among species of the two 100-nt intronic regions (5’ and 3’) that flank all human micro-exons and a set with the same number of randomly selected long exons. After training our CNN model, the micro-exon splicing event prediction accuracy, using an independent validation dataset, was 0.71 with an area under the ROC curve of 0.76, showing that our model had identified sequence patterns that have been conserved in evolution in the introns that flank micro-exons. Next, we introduced *in silico* point mutations at each of the 200 nucleotides in the introns that flank a micro-exon and used the trained CNN algorithm to predict splicing for every mutated intronic sequence version. This analysis identified thousands of point mutations in the flanking introns that significantly decreased the power of the CNN model to correctly predict a neighboring micro-exon splicing event, thus pointing to predictive bases in intronic regions important for micro-exon splicing signaling. We found these predictive bases to locate within conserved RNA-binding-motifs for RNA-binding-proteins (RBPs) known to relate to micro-exon splicing. Experimental data of minigene splicing reporter changes upon intron-base point-mutation confirmed the effect predicted by the CNN model for some of the micro-exon splicing events. The model can be used for validating novel micro-exons *de novo* assembled from RNA-seq data, and for an unbiased screening of introns, identifying genomic bases that have high micro-exon-splicing predictive power, possibly revealing critical point mutations that would be related in a yet unknown manner to a given disease.

## 1 Introduction

In eukaryotes, splicing events in pre-mRNAs from several mature transcripts culminate in the production of multiple protein isoforms produced from the same gene structure. These splicing events involve very precise and specific mechanisms which add another layer of complexity in gene regulation (Pan et al., 2008). About 90-95% of multi-exon genes are estimated to have alternative splicing isoforms, affecting the variability of expression between cells and tissues (Wang et al., 2008), and modifying cell localization and abundance of various protein isoforms that alter gene regulation of the cell (Gallego-Paez et al., 2017). In order to be a dynamic and well-orchestrated mechanism, several factors influence the splicing process such as spliceosome formation, involvement of RNA-binding proteins (RBPs) and participation of regulatory sequences such as intronic splicing enhancers/silencers (ISE/ISS) and exonic splicing enhancers/silencers (ESE/ESS), among others (Wang et al., 2015). Several studies have already pointed to altered splicing events in genes transcribed in cancer or neuropsychiatric diseases (Wang and Cooper, 2007; Suñé-Pou et al., 2017), for example in prostate cancer, where 30 % of the studied genes had only differences in their spliced isoforms and not in expression levels, showing the relevance of splicing regulation in functional biological processes. Thus, understanding the regulatory events in splicing can point out important aspects associated with the diseases.

Advances in large-scale technologies have pointed to a new class of exons, the so-called micro-exons, which were originally defined as exons which range in length up to 25 nucleotides (nt) (Volfovsky et al., 2003). Mammalian exons are on average 140-nt long (Gelfman et al., 2012), and the conventional splicing machinery has a predilection for exons with an average 140-nt length (Schwartz et al., 2009). Nevertheless, thousands of human genes have micro-exons (Li et al., 2015; Tapial et al., 2017), which are especially expressed in neuronal tissues at different stages (Yan et al., 2015). In invertebrates, we have shown that the *Schistosoma mansoni* parasite has over a dozen different micro-exon gene (MEG) families (DeMarco et al., 2010); each MEG has from 4 up to 19 micro-exons that generate protein variation through the alternate splicing of short (≤ 36 nt) symmetric exons organized in tandem (DeMarco et al., 2010). More recently, micro-exons were defined as exons with lengths ≤ 51 nt (Li et al., 2015). Given their short length, micro-exons would not accommodate large numbers of exonic splicing enhancers/silencers, requiring that these regulatory elements be primarily located in the introns that flank these micro-exons. Li *et al*. (Li et al., 2015) have shown that, in mammals, the conservation of bases in introns that flank micro-exons is greater than the conservation of introns that flank non-micro-exons. Another documented feature is the differential distribution of certain 6-base motifs (k-mers) in the intronic regions that flank micro-exons (Ustianenko et al., 2017). In addition, these short motifs are co-localized with some RNA binding proteins such as RBFOX and PTBP1, as evidenced by CLIP-seq assays with brain tissues and HeLa cells (Li et al., 2015). Silencing and overexpression assays for nSR100 protein showed a large effect on the mechanism of micro-exon splicing in 293T kidney cells (Irimia et al., 2014). In that study, Irimia *et al*. (Irimia et al., 2014) identified 126 micro-exon splicing events altered in the brain of autistic patients compared with controls, corresponding to 30 % of all micro-exon splicing events in that tissue.

The above set of information suggests that there might be specific mechanisms that define micro-exon splicing, but these mechanisms are still not fully explored. In fact, up until now most machine learning algorithms have searched for patterns in splicing events in general, such as SpliceAl (Jaganathan et al., 2019), not specifically looking for patterns involved with micro-exon splicing events.

Here, we performed a detailed computational search of patterns that could enable the splicing machinery to operate on micro-exons using a Convolution Neural Network (CNN) deep learning approach (Angermueller et al., 2016). More important, we have combined the CNN deep learning approach with an *in silico* point mutation strategy that scans the intronic sequences that flank micro-exons, in search for critically conserved bases where point mutations can be predicted to negatively impact the splicing of a neighboring micro-exon. Identification of such conserved intronic patterns involving micro-exon splicing extends the knowledge about the factors that control micro-exon splicing events in normal cells. It also opens the way for future large-scale screening of rare point mutations in the human genome that can change the intronic conserved patterns and would be predicted to impair processing of flanking micro-exons. Such an approach could accelerate the identification of intronic mutations that lead to micro-exon splicing defects yet unknown to be related with disease states.

## 2 Materials and Methods

### 2.1 Convolution Neural Network (CNN)

In order to train a classifier that could distinguish micro-exons (≤ 39 nt) from long exons (> 39 nt), all 4,908 micro-exons annotated in the human genome assembly (hg38) with the Ensembl annotation (GRch38.76) were identified; in order to have a balanced CNN model, an equal number of 4,908 randomly selected long exons was identified. For each selected exon, the 100-nt sequence from the intronic region upstream of the exon 5’-end and the 100-nt intronic sequence downstream of the 3’-end were extracted. To provide information to the classifier about conservation in the regions that flank micro-exons and long exons, the conservation score in vertebrates (PhastCon100way) for each nucleotide in the two 100 bp intronic regions that flank each of the selected exons was obtained.

The sequences *s* that flank each of the exons were transformed into categorical variables with the help of a *one-hot encoder* [A: (0,0,0,1) C: (0,0,1,0) T: (0,1,0,0) G: (1,0,0,0)] and the conservation values *c* were maintained as a continuous variable ranging from 0 to 1.

These data were used to train a Deep Convolutional Neural Network (CNN) with 1D convolutions from 4 different inputs. Inputs can be described as:

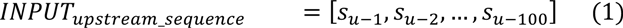

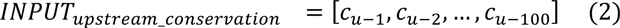

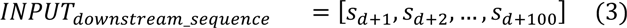

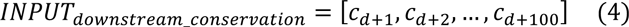

Where *u* represents the genomic coordinate of the 5’ end of an exon (either a micro-exon or a long exon) and *d* the genomic coordinate of the 3’end of the same exon. *s* and *c* represent respectively the vector containing the *one-hot encoder* and the nucleotide conservation value of a given coordinate in the flanking intron related to that exon.

The model was trained using binary crossentropy as the loss function, learning rate = 0.001, decay = 0.0 and optimized with rmsprop. The final dataset contained 7,067 exons for training (3,534 micro-exons and 3,533 long exons), and another 1,767 exons were used for validation (883 micro-exons and 884 long exons), while the remaining 982 sequences were used to assemble the ROC curves (491 micro-exons and 491 long exons).

Training was performed during 4000 epochs using a 500-size batch. The selected model was the one that obtained the highest accuracy in the validation data during the training. The analyzes were performed with keras (Chollet) in python 2.7.

### 2.2 *In silico* mutations

For each base within a 100-nt intronic region that flanks a micro-exon under analysis, with a given genomic coordinate, a *PositionScore* value was generated that corresponds to the importance of a given intronic nucleotide n for the prediction of the nearby micro-exon given the trained model. Calculation of the value per n position can be obtained as follows:

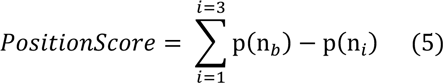

Where, variable *b* represents the nucleotide base found in the original sequence and variable *i* represents one of the 3 other possible nucleotides. The p function is the prediction value of the micro-exon by the trained CNN model given all the original parameters of the intron that flanks a micro-exon, or after nucleotide n*_b_* is replaced by nucleotide n*_i_* at the given position. The final *PositionScore* value of a given n position in any intron that flanks a micro-exon was determined by the sum of the differences between the original base prediction value and the artificially mutated base prediction values.

All possible positions (100 upstream and 100 downstream) in the introns that flank all 4,908 micro-exons had their *PositionScore* calculated. After that, the *PositionScores* with negative and positive values were normalized by the *PositionScore* with the lowest, most negative and the highest, most positive values, respectively, sorted from the lowest, most negative to the highest, most positive score, and the genomic coordinates of the positions having the top 5 % *PositionScores* with the most negative values were annotated.

### 2.3 Base Group Evaluation

The top 5 % intron bases that had the greatest negative influence on micro-exon prediction (most negative *PositionScore* values) were divided into five different groups, each group having the same number of intron bases but with different *PositionScore* values. GroupA represents the top 1 % quantile with the most negative *PositionScore* values and GroupE the lowest of the top five 1 % quantiles. The distance measurements between the intron bases and the 5’-end and 3’-end of the exon were obtained using BEDTools (v2.26.0) (Quinlan and Hall, 2010), and comparisons of the distributions were performed with Kolmogorov-Smirnov test. Density distributions of distances from bases to the 5’- or 3’-end were obtained with ggplot2. Kendall’s rank correlation test was used to obtain the correlation between the absolute *PositionScore* values of the bases and the distances. For interspecies conservation all PhastCon data (Siepel et al., 2005; Pollard et al., 2010) for primates, placentals, and vertebrates were used (PhastCon7way, and PhastCon100way). Conservation scores were compared using the Kolmogorov-Smirnov (KS) test calculated with R (R Core Development Team, 2013).

### 2.4 Motif analyses

In silico analyses of the RNA-binding-motifs were performed using the MEME program (v. 4.12.0) (Bailey et al., 2009). All bases in the intronic regions that were identified by the machine learning (ML) algorithm as having negative influence on micro-exon prediction were extended by 5 nt at both ends (5’ and 3’), resulting in an 11-nt-long sequence; for this, the respective genomic sequences were retrieved from ENSEMBL (hg38) assembly (ftp://ftp.ensembl.org/pub/release-96/fasta/homo_sapiens/dna/Homo_sapiens.GRCh_38.dna_sm.primary_assembly.fa.gz) using getfasta from BEDTools tool (v2.26.0) (Quinlan and Hall, 2010) according to the strand orientation of the transcript. MEME (Bailey et al., 2006) was used to identify the 3 most hyper-represented 11-nt-long sequences in each of the five groups (GroupA to GroupE, see above) in a non-biased way using a zero-order probability model; only single nucleotide frequencies would be measured without di- or tri-nucleotides. To build the zero-order probability model, the 100-nt-long upstream and downstream intronic sequences that flank long exons (> 39 nt) (n = 4,417 exons, 8,834 intronic flanking sequences, model: A = 0.246, C = 0.215, G = 0.222, T = 0.317) were scanned using the 11-nt-long sequences. The parameters used for this analysis were: zoops (Zero or One Occurrence Per Sequence), number of motifs identified = 3, and the window size representing the motif size was 6 to 8 nt. Only motifs with E-value ≤ 0.05 were considered for further analyses.

### 2.5 Motif identification

Motif identification was performed using the TomTom similarity algorithm (Gupta et al., 2007). The enriched motifs identified in the previous analysis were used as query sequences and the targets were the sequences that are deposited in the ATtRACT Database (Giudice et al., 2016). This database compiles information on 370 RBPs and 1583 RBP consensus RNA-binding-motifs; only human genome sequences were used, resulting in 1,094 consensus sequences. The similarity matrix used was the Euclidean distance, which has a higher accuracy rate when compared with other functions (Gupta et al., 2007). Only sequences that had similarities with E-value ≤ 0.05 were selected for further analysis. Sequence data representing intronic splicing enhancers / silencers (ISE/ISS) were used as targets in an additional search for similarity. Intronic Splicing Enhancers (ISE) sequences were obtained from the study by Wang et al. (Wang et al., 2012) represented by 109 sequences. To reduce redundancy only the main clusters were used, representing six consensus sequences (GGGTTT, GGTGGT, TTTGGG, GAGGGG, GGTATT and GTAACG). Sequences referring to ISS (Intronic Splicing Silencers) were obtained from the study by Wang et al. (Wang et al., 2013b) represented by 102 sequences. The same strategy was used to avoid redundancy, and only the main groupings were used as the target sequence, resulting in 10 consensus sequences (CACACCA, CTCCTC, UACAGCT, CTTCAG, GAACAG, CAAAGGA, AGATATT, ACATGA, AATTTA and AGTAGG).

### 2.6 Motif enrichment

Motif enrichment analysis was performed using the CentriMo algorithm (Bailey and MacHanick, 2012). Only data from the RBP sites identified in the similarity analysis (Motif identification, see above) was used in each analysis to reduce the multiple testing rate. As a negative control, sequences of intronic regions (100 nt) from the long exon model were used to calculate enrichment in a 6 to 8 nt window, with all other default parameters. First, the algorithm uses a 6- to 8-nt window to identify motifs along the given intronic sequence and calculate the significance of enrichment at a specific location, given by a p-value, which was corrected for multiple testing and represented by E-value. After this step, the frequency of similar sequences in the data of interest was calculated and compared with the negative control sequences, the significance of the difference between expected and observed was given by the result of the Fischer test adjusted for multiple testing. To perform this analysis, the sequences were divided into upstream and downstream from exonic and micro-exonic regions, due to the fact that some binding sites were enriched in upstream regions and not downstream and vice versa.

### 2.7 Gene Ontology (GO) enrichment analysis

GO enrichment analysis was performed with Webgestalt (Wang et al., 2013a) using over representation analysis (ORA) with at least 3 genes and as background all cataloged human proteins. False Discovery Rate (FDR) ≤ 0.05.

### 2.8 eCLIP assay data

eCLIP-seq data was downloaded from the ENCODE project portal (Davis et al., 2018) at (https://www.encodeproject.org). Data from K562 and HepG2 cell lines, for **PTBP1** (ENCFF051PIE, ENCFF245YUN, ENCFF363UDO, ENCFF936SHU, ENCFF476HFB, ENCFF556EQK, ENCFF258TKH, ENCFF617YCT, ENCFF799AHI, ENCFF967LWB, ENCFF207EDD, ENCFF665CYG), **TIA1** (ENCFF093IND, ENCFF873ZAY, ENCFF782ZMF, ENCFF940BFP, ENCFF951BGZ, ENCFF573VNX, ENCFF996GFV, ENCFF306MBI, ENCFF048JJS, ENCFF625OCH, ENCFF523SWX, ENCFF698IQD) and **U2AF2** (ENCFF368XEI, ENCFF159SPZ, ENCFF536AFD, ENCFF913WRH, ENCFF566CFJ, ENCFF989JBA, ENCFF765TAB, ENCFF712LBW, ENCFF524JHH, ENCFF024JFG, ENCFF945AJC, ENCFF126CZT) RBPs were used. Data for eCLIP-seq (Van Nostrand et al., 2016) in bigWig format was obtained, both for the target proteins of interest and their respective controls (mock IgG). The files were converted to wig using the UCSC tools (Kent et al., 2010). The raw signal from wig files were represented by signal+1 in order to be more stringent to small values and avoid 0 division. The median signal of each assay was calculated for each of the groups and for the long exons negative control and divided by the signal of the respective mock control.

### 2.9 RBP knock down (KD)

RNA-seq data in HepG2 and K562 cells for knock down of *PTBP1* (ENCFF001ZGD, ENCFF001ZGF, ENCFF001ZGI, ENCFF001ZGJ, ENCFF184CDV, ENCFF456OPJ, ENCFF486ADH, ENCFF555EDL, ENCFF642KBO, ENCFF887GOE, ENCFF893AGN, ENCFF983TGB), for knock down of *U2AF2* (ENCFF158ZML, ENCFF593VXV, ENCFF550GXB, ENCFF424URS, ENCFF470BBN, ENCFF235FRZ, ENCFF026PLZ, ENCFF824GIZ, ENCFF020XNK, ENCFF229BQW, ENCFF298TSM, ENCFF354AMD), for knock down of *TIA1* (ENCFF741EQA, ENCFF578TWY, ENCFF695YNR, ENCFF338TUE, ENCFF773CAF, ENCFF647VRD, ENCFF228TQK, ENCFF845KED) and for controls (ENCFF385GEX, ENCFF403CZA, ENCFF278TEH, ENCFF922CDR, ENCFF910EGI, ENCFF430ZBY, ENCFF291QQH, ENCFF503VRZ, ENCFF105YHI, ENCFF602GIQ) were obtained from the ENCODE project portal (Davis et al., 2018) at (https://www.encodeproject.org), and the data is described in (Nostrand et al., 2018). Reads were quality checked with FastQC (v0.11.7) (Andrews, 2010) and the adapters removed with Fastp (v0.20.0) (Chen et al., 2018). Reads mapping to exon splice junctions and differential abundance analyses were performed using vast-tools (v2.2.2) (https://github.com/vastgroup/vast-tools#vast-tools-1) and the human genome assembly (hg19). The database used contains 402,157 reference splicing events described in the Vertebrate Alternative Splicing and Transcription Database (VastDB) (Tapial et al., 2017) for the human genome; VastDB is one of the largest resource of genome-wide, quantitative profiles of AS events assembled to date. The vast-tools use Bowtie (Langmead, 2010) for genome mapping; first, reads were divided into 50-nt over a 25-nt window, after this process the reads that have not been mapped to the genome were used for mapping at known exon splice junctions, and for *de novo* junctions a model was built where the 5’- and 3’-end of the same exon needed to be less than 300 nt apart and the junction must have had the canonical splice site donor/acceptor GU/AG (Irimia et al., 2014).

### 2.10 SDVs dataset

The Multiplexed Functional Assay of Splicing using Sort-seq (MFASS) dataset that has been generated to determine splice-disrupting variants (SDVs) was downloaded from the work by Cheung *et al* (Cheung et al., 2019) and compared with the list of introns that flank micro-exons identified in our CNN model. There were 27,733 rare variants from the ExAC database assayed by MFASS and 1,050 classified as SDVs (Cheung et al., 2019). Comparisons were performed using BEDTools tool (v2.26.0) (Quinlan and Hall, 2010) with the intersect function with -f 1 -r parameters. All comparisons were performed using hg38 assembly coordinates.

## 3 Results

### 3.1 Prediction of splicing of micro-exons with the Convolutional Neural Network algorithm using primary sequence and conservation score among vertebrates

In order to determine whether the pattern of bases conservation in introns interferes with micro-exon splicing events in humans, we constructed a prediction deep learning model using a Convolutional Neural Network (CNN) (Figure 1A), which was based on both the sequence content of the 100-nt-long intronic regions that flank micro-exons and long exons at their 5’- and 3’-ends, and the sequence conservation among the species of these 100-nt-long intronic sequences (Figure 1A). Conservation score values for the human genome bases obtained by comparison with 100 vertebrate genomic sequences (Siepel et al., 2005, 2006) were used to obtain the conservation level of intronic regions that flank the micro-exons and long exons (+100 bases downstream to exons and −100 bases upstream to exons).

**Figure 1.**
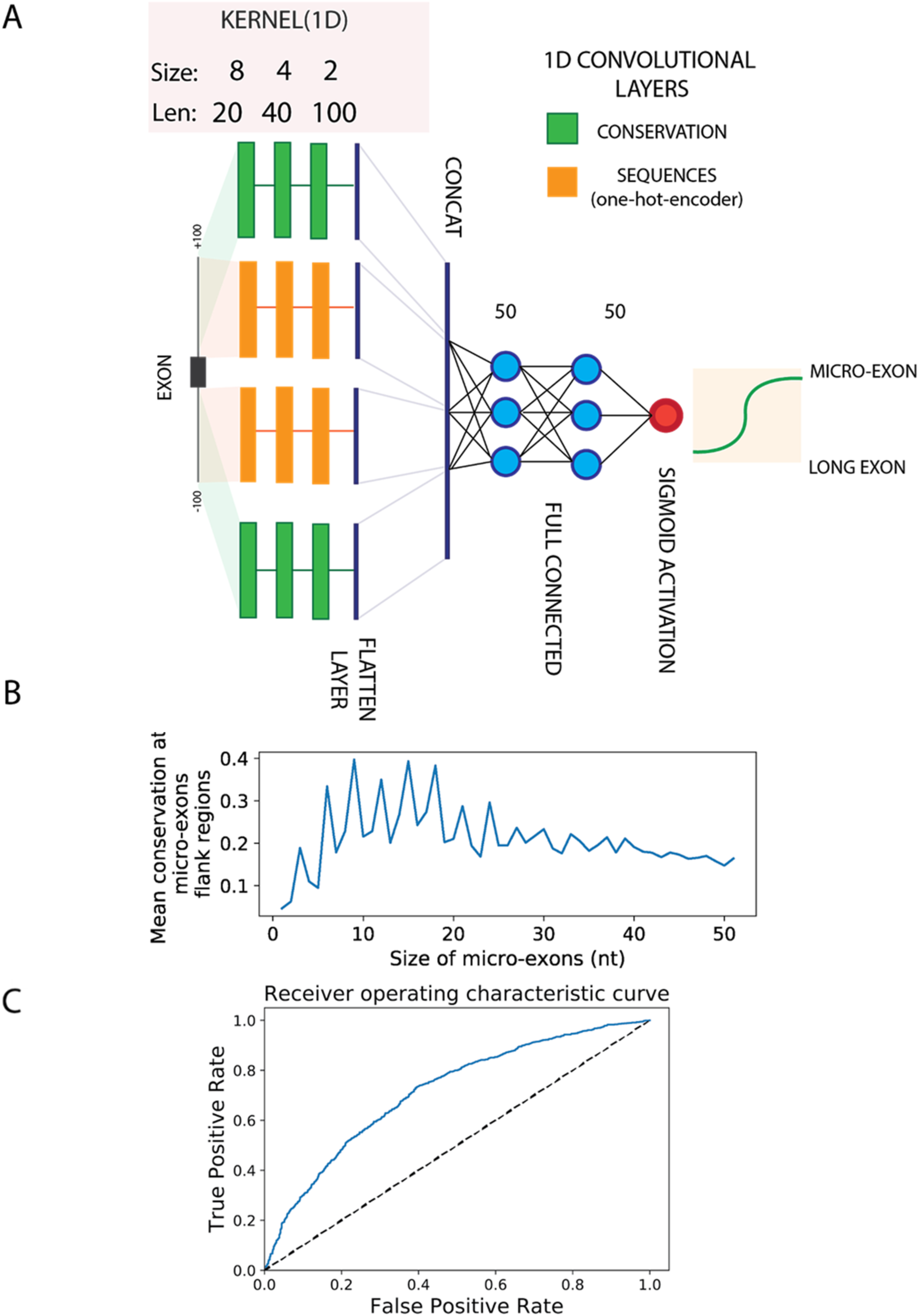
Deep Learning CNN scheme and identification of conserved sequence patterns in the introns that flank micro-exons. **(A)** Deep Learning Convolutional Neural Network (CNN) architecture used for the classification of micro-exons and long exons based on the sequences of their flanking intronic regions and on the interspecies conservation of these introns. This neural network architecture consists of separate input data of the intronic sequences that flank the exons (orange) and of interspecies conservation data for these introns (green), for both the downstream and upstream 100-nt regions that flank the exons. Flanking sequences were processed in two input convolution layers that turn the sequences into recurring motifs in a succession of 3 convolutional layers with different kernel lengths. Input data containing the conservation level of the sequences were convoluted separately using 3 other convolutional layers with the same kernel number and length of the flanking sequence layers. The output of the 4 convolutional layers were flattened and then concatenated in a unique layer (CONCAT) containing the motif-convolved sequences. This data was propagated through two sequential fully-connected layers (blue) that output a binary classifier (red dot) containing a sigmoidal function that can discriminate a flanking micro-exon from a long exon. **(B)** Mean conservation (y-axis), calculated by the CNN model, of the first upstream and downstream 100 nucleotides that flank micro-exons of different lengths (in number of bases), as indicated on the x-axis. **(C)** Receiver Operator Characteristics (ROC) curve for the CNN model classification of exons based on the conservation pattern of their 5’- and 3’-flanking introns. The Area Under the Curve (AUC) was 0.76 for the prediction performed with an independent validation dataset, using the intronic sequences that flank micro-exons (≤ 39 nt) and long exons (> 39 nt). The dotted line represents the accuracy values for a random model (AUC = 0.5).

These values were then used to assess the conservation of introns that flank micro-exons of different lengths. We observed that introns that flank symmetrical micro-exons (micro-exons whose lengths were an exact multiple of 3) were more conserved than introns that flank non-symmetrical micro-exons (Figure 1B, see peaks at 3, 6, 9 nt, etc.). This difference in intronic conservation as a function of micro-exon length was no longer noted for introns that flank micro-exons over 39 nt in length (Figure 1B). Using the conservation information of Figure 1B, we decided to train our CNN model (Figure 1A) using introns that flank micro-exons only of lengths ≤ 39 nt. This choice assumes that the elements that are recognized by the splicing machinery were conserved during evolution in the intronic regions that flank the micro-exons.

To train the CNN model, we retrieved all 4,908 micro-exons of lengths ≤ 39 nt that were present in the Ensembl annotation (GRch38.76) of the hg38 version of the human genome, and randomly divided the set in three parts: 10 % (491 micro-exons) were set aside for the final test of performance of the model; of the remaining 4,417 micro-exons, 80 % were used for training the model (3,534 micro-exons), while 20 % (883 micro-exons) were used for an independent validation of the trained model. In order to have a balanced CNN model, an equal number of 4,908 randomly selected long exons (> 39 nt) was used.

The CNN model was trained with the set of 7,067 intronic 100-nt-long sequences that flanked the 3,534 micro-exons, both upstream of the micro-exon 5’-end and downstream of the 3’-end. An equal amount of 7,067 intronic 100-nt-long sequences that flanked 3,534 long exons on both ends was also used for training. With the trained CNN model, a micro-exon prediction accuracy of 0.71 was obtained for a validation test with an independent dataset of 1,766 intronic regions, and the area under the ROC curve was 0.76 (Figure 1C).

As a parallel control, we tested the performance of the CNN model only using the intronic sequences, without the conservation scores; after training without the conservation, the best obtained micro-exon prediction accuracy was only 0.59 for the validation test with the independent dataset, and the area under the ROC curve was 0.61. Given the low performance of this sequence-only model, we did not explore it further.

The observation that a good prediction accuracy was obtained with the complete CNN model, using both the sequences and their conservation scores, reinforces the idea that our machine learning approach was finding a pattern in flanking introns that has been conserved in evolution, and that should participate in micro-exon processing events.

### 3.2 *In silico* point mutations in the introns that flank micro-exons affected the splicing predictive power of the CNN model

Next, we introduced *in silico* point mutations, one at a time, at each of the 200 nucleotides that flank the micro-exons or long exons, replacing the original base with each of the 3 other bases, and the CNN trained algorithm was used to predict splicing for every mutated intronic sequence version. The objective of this strategy was to estimate to what extent the conservation at each position along the intron interfered with the CNN model classification of the nearby micro-exon or long exon splicing event. The difference in the predictive value obtained before and after the *in silico* point mutation of each base was summarized with the *PositionScore* value for the respective base, as described in the Methods.

A heatmap of *PositionScore* values along the introns that flank all tested micro-exons was generated (Figure 2), with the micro-exons being clustered according to the pattern of *PositionScores* across their flanking introns. The heatmap shows that when each original base was *in silico* mutated, being replaced by each of the other 3 bases, negative and positive *PositionScore* values were obtained for many of the bases along the introns that flank each micro-exon (Figure 2). This indicates that the micro-exon-splicing prediction power of the CNN model was altered by the resulting *in silico* mutated intron, compared with the original intron sequence. Bases with negative *PositionScores* (Figure 2, blue regions) indicate that a given point mutation had a negative impact on the prediction power, i.e. the mutation resulted in an increased likelihood that the intronic sequence were mistakenly classified by the model to be an intron that flanks a long exon (Figure 2). These data highlight conserved sequences in the flanking introns important for micro-exon splicing signaling. The red points in the map (Figure 2) show that when *in silico* point mutations were introduced at certain points in the introns, there was an increased likelihood of those sequences being recognized by the CNN model as introns that flank micro-exons. This could be due to the fact that, for that given intron, the power of the CNN model classification might have been near the significance cutoff when the wild-type sequence was considered, while in the *in silico* mutated sequence the change in a base in the red region has possibly changed its sequence pattern towards a more robust, conserved pattern of introns that flank micro-exons.

**Figure 2.**
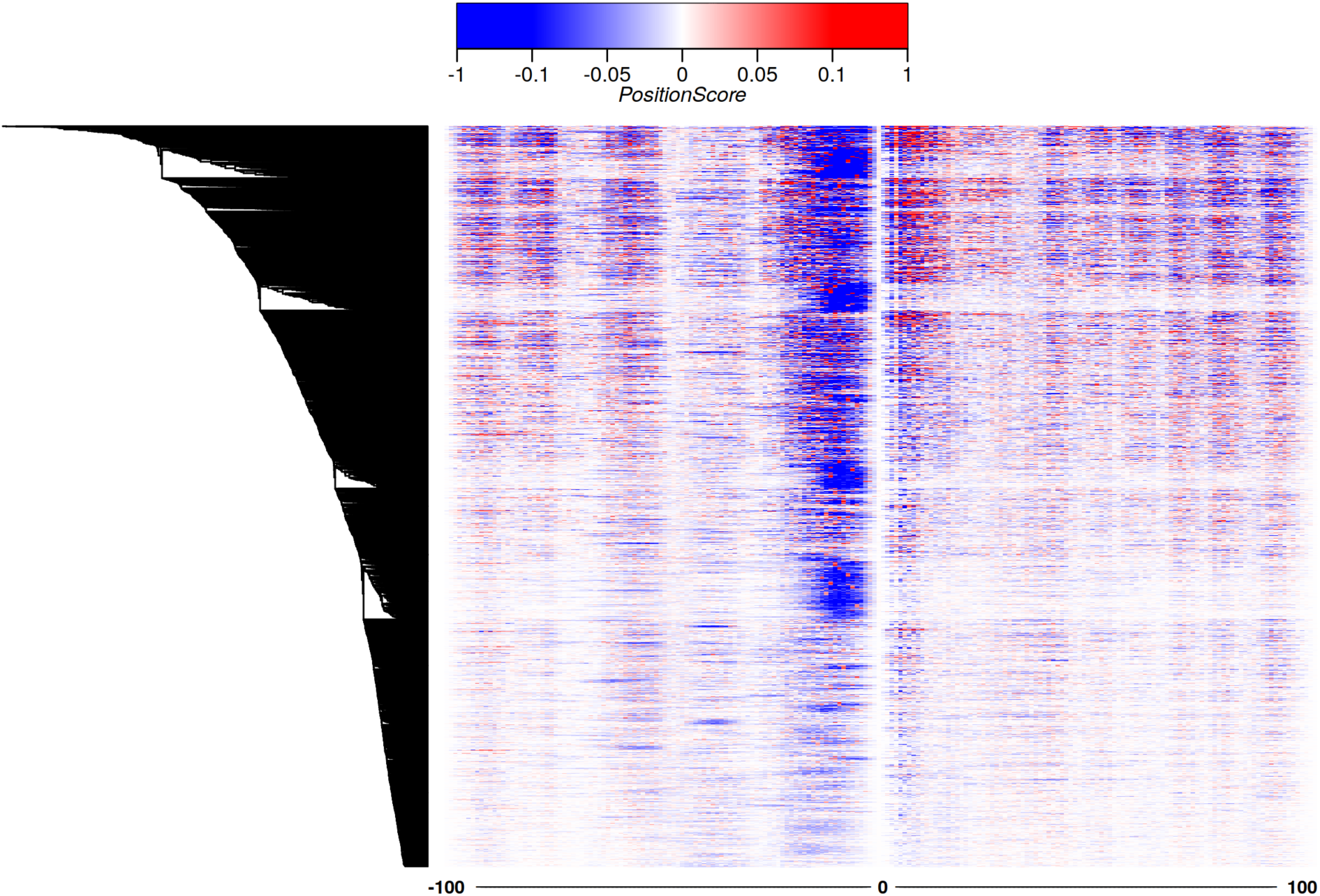
Positions of critical bases conservation along the introns that flank all human micro-exons. On the y-axis each of the 4,417 micro-exons, that were used in training and validation of the Deep Learning CNN model, is represented in one line. The x-axis shows the 200 nucleotides that flank each micro-exon (100 nt at the 5’ or 3’ ends); for each base of the intron sequence that flanks the micro-exon, the delta value (*PositionScore*) of the prediction perturbation caused by the in silico point mutation of that base is represented by the color; the delta was calculated by subtracting the intronic sequence prediction value, obtained after the base at a given position was changed to the other 3 possible bases, from the intronic sequence prediction value using the original wild-type base. The heatmap has clustered the micro-exons according to the *PositionScore* pattern of the intron sequences that flank each micro-exon. On the upper left is the color scale of the perturbation *PositionScore* values. Positive values indicate that the *in silico* mutation increased the probability that a given sequence was classified as an intron that flank a micro-exon, and negative values show that the in silico mutation increased the probability that the sequence was mistakenly classified as an intron that flank a long exon.

### 3.3 Different density distribution of predictive *PositionScore* values for bases in the introns that flank micro-exons

To show that the machine learning approach was pointing to a sequence pattern conserved during evolution, the *PositionScores* were compared with PhastCon conservation scores across different species. First, the predictive bases were grouped according to the value of the *PositionScore*, in the following way. Bases were ordered according to the values of *PositionScore*, from the lowest, most negative to the highest, most positive. Bases with the lowest, most negative *PositionScores* represent nucleotides with the greatest negative impact on micro-exon-splicing predictive value. In total, 23,704 bases were pooled, originating from analysis of the introns that flank all 4,417 micro-exons; these 23,704 bases represent the top 5 % with the lowest, most negative *PositionScores* (out of the 474,079 bases with *PositionScores* < 0). The 23,704 bases were divided into five groups, with GroupA containing the bases with the lowest, most negative *PositionScore* values representing the top 1 % of the total bases (n = 4,741), and each of the four remaining groups were comprised of n = 4,741 bases (Table 1). The distribution of mean values of absolute *PositionScore* along the five groups from GroupA to GroupE is shown in Supplementary Figure 1A. The difference in the median absolute *PositionScore* between GroupA and GroupB was the largest (0.14) (Table 1).

**Table 1.**
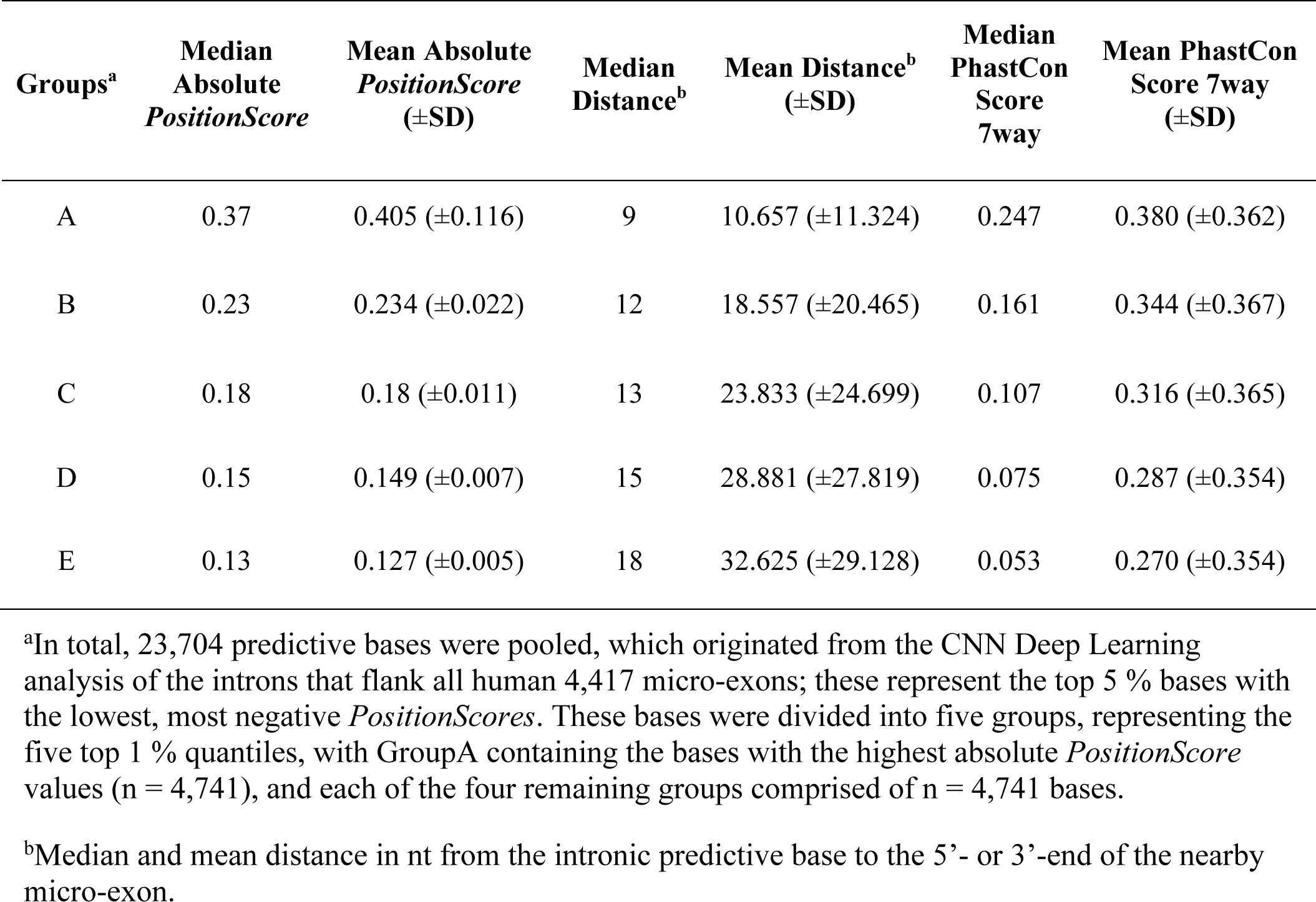
Groups of introns that flank micro-exons and have different micro-exon-splicing predictive *PositionScore* values

| Groups<sup>a</sup> | Median Absolute *PositionScore* | Mean Absolute *PositionScore* (±SD) | Median Distance<sup>b</sup> | Mean Distance<sup>b</sup> (±SD) | Median PhastCon Score 7way | Mean PhastCon Score 7way (±SD) |
| --- | --- | --- | --- | --- | --- | --- |
| A | 0.37 | 0.405 (±0.116) | 9 | 10.657 (±11.324) | 0.247 | 0.380 (±0.362) |
| B | 0.23 | 0.234 (±0.022) | 12 | 18.557 (±20.465) | 0.161 | 0.344 (±0.367) |
| C | 0.18 | 0.18 (±0.011) | 13 | 23.833 (±24.699) | 0.107 | 0.316 (±0.365) |
| D | 0.15 | 0.149 (±0.007) | 15 | 28.881 (±27.819) | 0.075 | 0.287 (±0.354) |
| E | 0.13 | 0.127 (±0.005) | 18 | 32.625 (±29.128) | 0.053 | 0.270 (±0.354) |

<sup>a</sup>In total, 23,704 predictive bases were pooled, which originated from the CNN Deep Learning analysis of the introns that flank all human 4,417 micro-exons; these represent the top 5 % bases with the lowest, most negative *PositionScores*. These bases were divided into five groups, representing the five top 1 % quantiles, with GroupA containing the bases with the highest absolute *PositionScore* values (n = 4,741), and each of the four remaining groups comprised of n = 4,741 bases.
<sup>b</sup>Median and mean distance in nt from the intronic predictive base to the 5'- or 3'-end of the nearby micro-exon.

Analysis of the distribution of GroupA predictive base positions along the introns that flank the micro-exons showed predictive bases more densely located at a median distance of 9 nt up- and downstream from the micro-exon ends (Figure 3A, Table 1), whereas in GroupB the median was 12 nt (Table 1). The average absolute values of *PositionScores* as a function of the distance to the micro-exon end was computed in 20-nt-long windows along the intron (for all groups A to E together) (Supplementary Figure 1B). As the distance between predictive base and micro-exon end increases, the absolute values of *PositionScore* along the intron decrease (Kendall’s rank correlation, tau = −0.23, p-value < 2.2e-16, Supplementary Figure 1B). All comparisons between groups showed a difference in distribution as a function of distance (Supplementary Table S1, Komogorov-Smirnov test, p-value < 0.05).

**Figure 3.**
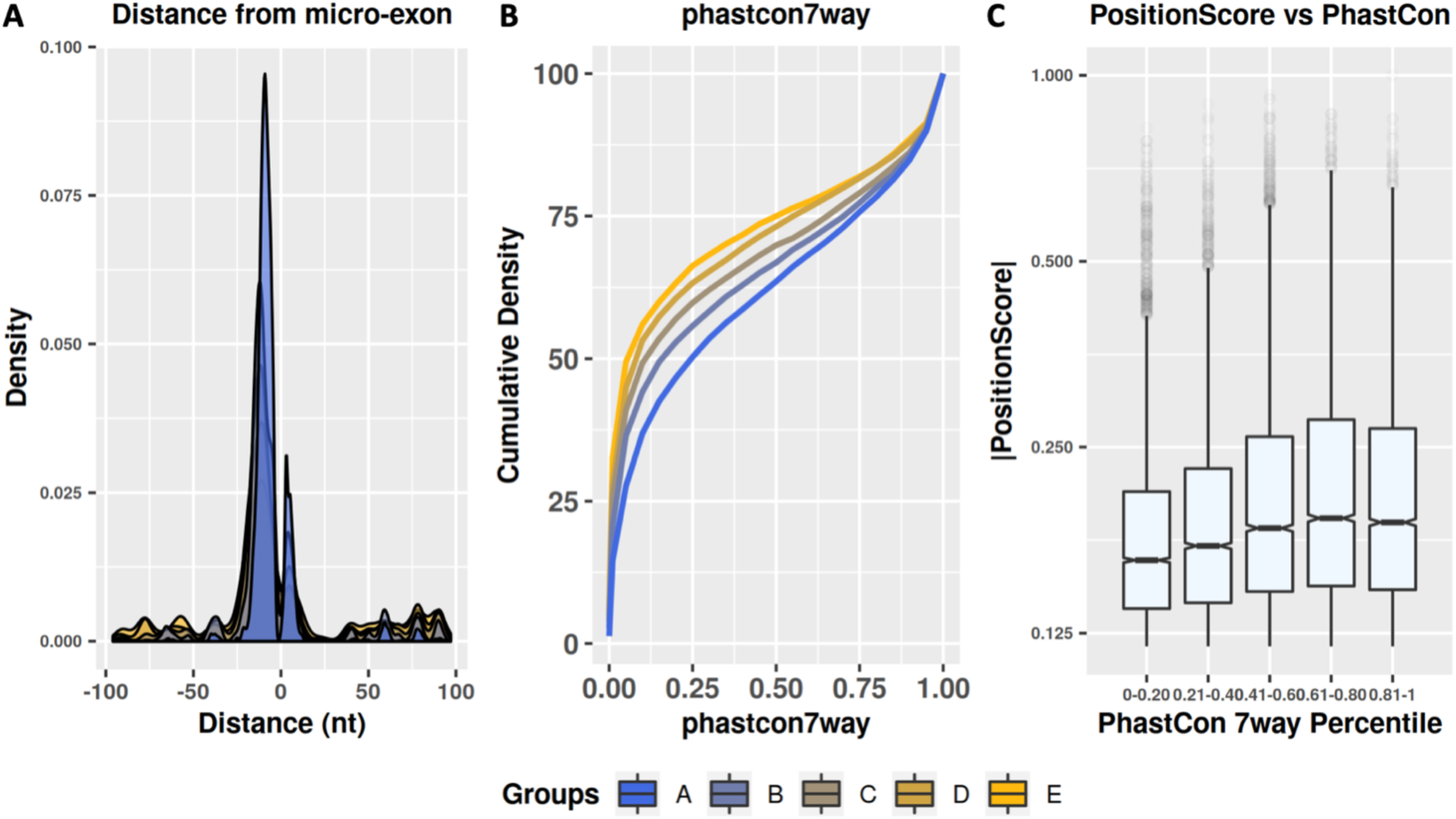
Splicing-predictive bases *PositionScore* distribution along the introns that flank micro-exons. **(A)** Density distribution of intron predictive bases (y-axis) as a function of distance to the micro-exons (x-axis), either upstream (−1 to −100 nt) or downstream (+1 to +100 nt) of the micro-exon ends. Distance equal to 0 marks the micro-exon. Each group is shown with a different color, as indicated at the bottom. Comparison of distribution between groups showed statistically significant difference (Kolmogorov-Smirnov test, p-value < 0.05), except for GroupD vs GroupE. **(B)** Cumulative distribution of PhastCon7way values for each of the five groups, indicated by the colors. The y-axis shows the cumulative distribution and the x-axis shows the PhastCon7way score. Statistical differences in PhastCon7way scores distribution were observed in all comparisons using GroupA or GroupB (Kolmogorov-Smirnov test, p value <0.05). **(C)** Box plot of absolute values of *PositionScore* of intron predictive bases (y-axis) as a function of PhastCon7wayScore computed in intervals of 20 percentile (x-axis). All *PositionScores* from the five groups (GroupA to GroupE) were plotted together. Correlation between *PositionScore* and PhastCon7wayScore was calculated (Spearman’s correlation, rho = −0.14, p-value < 0.05).

Since GroupA showed predictive bases closer to the micro-exons and larger absolute values of *PositionScore*, these bases were expected to be in more evolutionarily conserved regions compared with the other groups. As expected, in the analysis comparing the PhastCon7way values, which represent the conservation values among 7 vertebrates, GroupA showed higher conservation values when compared with the other groups; the cumulative density of the PhastCon7way value for each group shifted to the left as the group mean absolute *PositionScore* value decreased (Figure 3B). For GroupA bases the median value of PhastCon7way was 0.247, while in GroupB it was 0.161 (Table 1). All comparisons of PhastCon7way value distributions showed statistical difference (KS test, p-value < 0.05). The same pattern was observed when other PhastCon Scores background conservation values for 100 species were used (Supplementary Figure 2 and Supplementary Table S2). A statistically significant low correlation was observed between *PositionScore* and PhastCon7way (Spearman Correlation rho = −0.14, p-value < 0.05); it can be seen that higher PhastCon7way Scores were associated with higher absolute *PositionScore* values (Figure 3C).

**Table 2.**
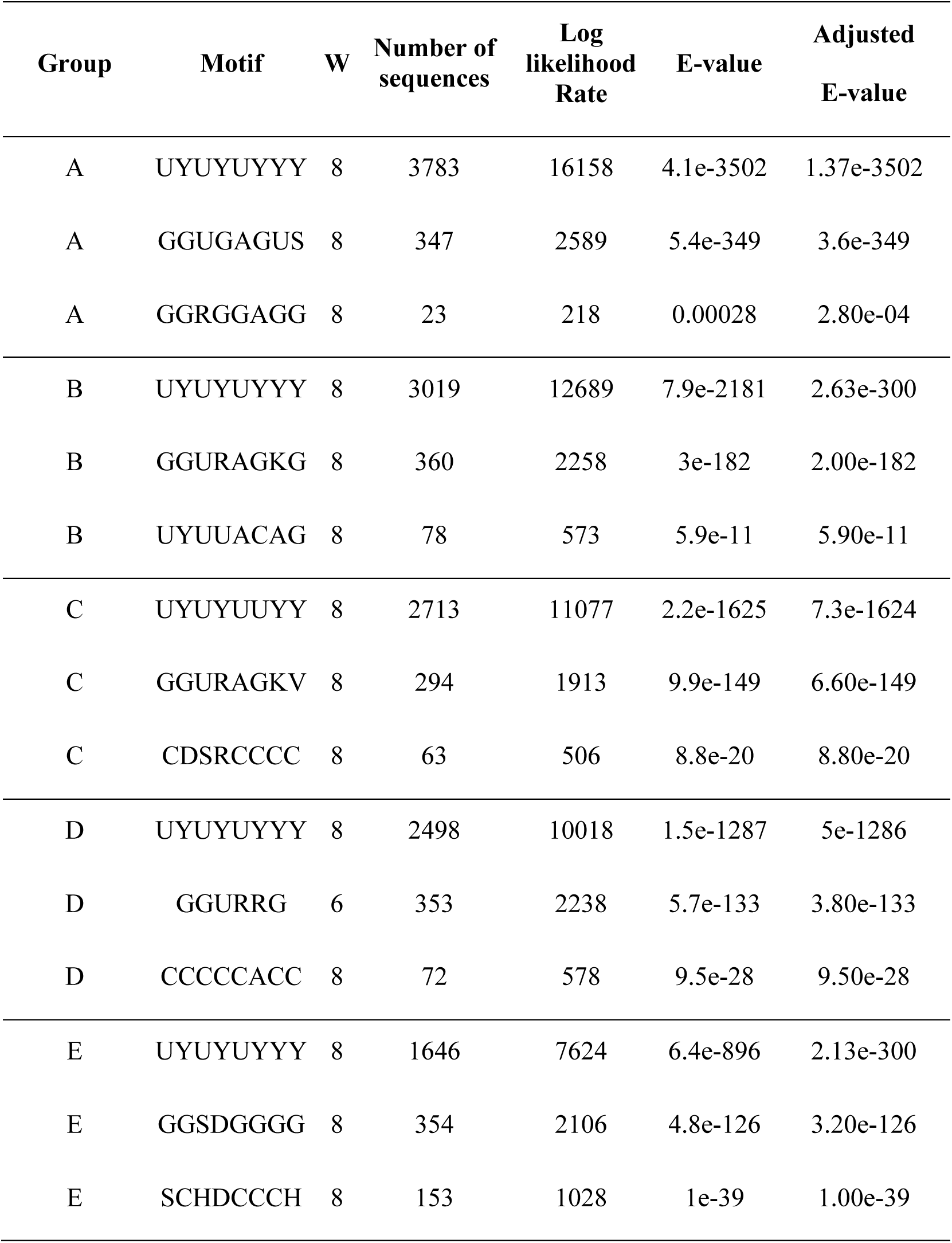
Enriched motifs in the introns that flank micro-exons (≤ 39 nt)

### 3.4 Identification of enriched specific sequences shows that the CNN model highlighted a homogeneous sequence pattern of predictive bases

To identify possible enriched sequence patterns containing the predictive bases (with the highest absolute *PositionScores*) within each group, the MEME Suite algorithm (Bailey et al., 2009) was used. For this purpose, a window was created with the five nucleotides present in the intron genomic sequence on each side of the predictive base under study, generating a small 11-nt-long sequence containing the predictive base. To identify over-represented sequences in each group, we contrasted the frequency of 11-nt-long intron sequences of GroupA with a background model using the base frequency of 11-nt-long windows along the 100-nt-long up- and downstream intronic regions that flank long exons.

In this analysis, the algorithm sought, within the 11-nucleotide sequences, to obtain a multiple alignment of all sequences from the same group, with at least 6 to 8 nucleotides aligned in each sequence, to ensure that the predictive base was included within the RNA motif to be found. The results shown in Table 2 include the 3 most abundant motifs in each group. It is worth noting that GroupA had more sequences that matched each of the 3 consensus motifs, suggesting that the bases with the highest absolute *PositionScores* housed more defined patterns. For example, in GroupA the number of sequences that aligned to generate Motif 1 (UYUYUYYY) was 3,783 (out of 4,741 sequences, 80%), while GroupB Motif 1 (UYUYUYYY) had 3,019 sequences (64 %), and the number of sequences within the enriched motifs decreased as a function of the lowering of the base predictive value in the groups (Figure 4A).

**Figure 4.**
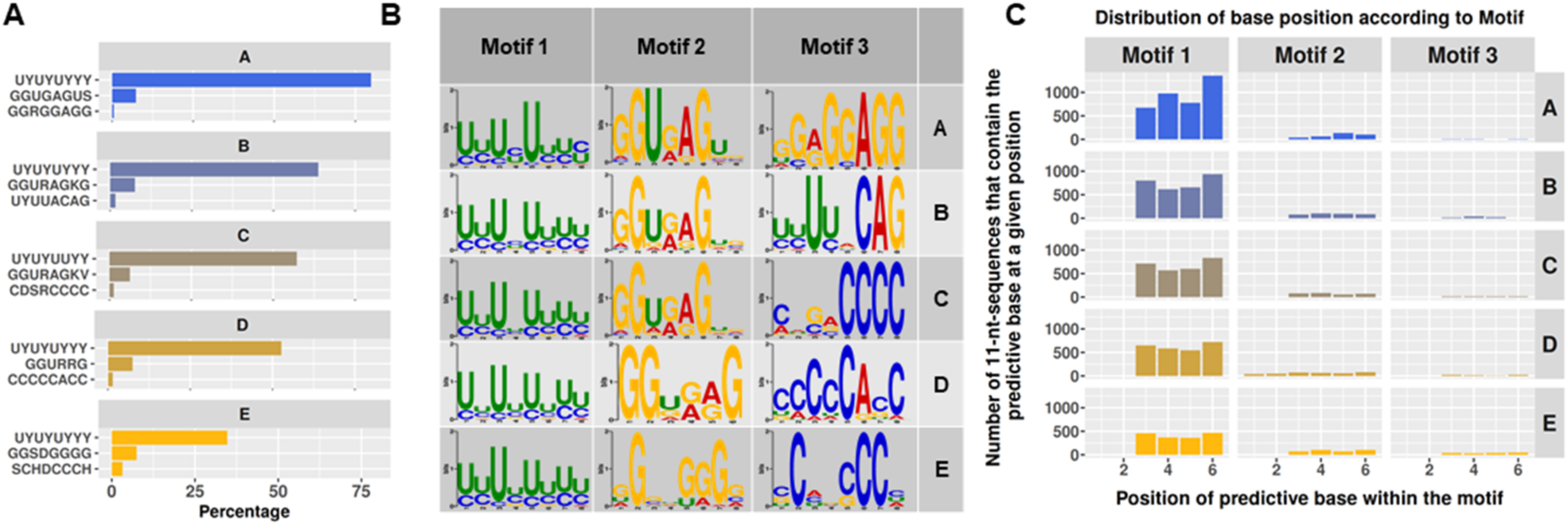
Enriched sequence motifs that contain the predictive bases identified by the CNN algorithm in the introns of neighboring micro-exons. **(A)** Number of intron sequences that support the enrichment of each motif indicated at left. The x-axis shows the number of intron sequences that contain the predictive bases within the corresponding motif, calculated as a percentage of the total number of sequences in the corresponding group (A to E). The y-axis shows for each group the sequence motifs 1 to 3, ordered from top to bottom according to the enrichment significance (based on the adjusted E-value). **(B)** Panels with logo sequences of the conserved motifs 1 to 3 in each group A to E, indicated at right. **(C)** Distribution of the relative position of the predictive bases within each motif, for motifs 1 to 3. Predictive bases were determined by the Deep Learning CNN algorithm as important in the intron for predicting neighboring micro-exon processing, in each group (A to E, as indicated at right). The x-axis shows the position within the enriched motif and the y-axis shows the number of predictive bases that were located at the corresponding position.

The two most enriched motifs were very similar among the groups, being comprised of sequences with high C and U contents or having a G-rich region. The third most enriched motif was characterized by the presence of a high-C content (Figure 4B). The frequency of predictive base position within the motif was different among sequences in the same group, and also different when comparing motifs between groups (Figure 4C), although the identified motifs were very similar among groups. Thus, Motif 1 in GroupA (Figure 4C) had more predictive bases located at positions 4 and 6, and in GroupB at positions 3 and 6, while in Groups C, D and E, the predictive bases were located mainly at the sixth position (χ^²^ test, df (12), p-value < 2e-16). Motif 2 in GroupA (Figure 4C) had more bases located at positions 5 and 6, whereas in GroupB at positions 4 and 5 and in GroupC at positions 3 and 4 (Figure 4C). GroupD predictive bases (Figure 4C) were located more frequently at position 6, and in GroupE at positions 3 and 6 (χ^²^ test, df (20), p < 2e-16). Motif 3 showed predictive bases widespread among positions 3 to 6 (Figure 4C), mostly at position 4 for Groups A, B and C, position 3 for Group D and position 6 for Group E (χ^²^ test, df (12), p = 4e-6).

### 3.5 Enriched motifs containing the predictive bases identified by CNN were enriched in RNA-binding-motifs of RBPs involved with RNA splicing

To test whether the motifs containing the predictive bases were similar to known RBP RNA-binding-motifs, we used the TomTom algorithm (Gupta et al., 2007) and the sequences were compared with the ATtRACT database (Giudice et al., 2016). This database is comprised of canonical and non-canonical RNA consensus sequences that are known binding targets of human RBPs. Searching for the three most enriched motifs that contained the predictive bases in all groups (A through E), six RBPs were found in common among the analyzes (Figure 5A, Table 3), namely PTBP1, ELAVL1, U2AF2, ELAVL2, TIA1 and PCBP1. The PTBP1 (Polypyrimidine Tract Binding Protein 1) motif was detected in all groups with the highest significance score. GO biological process enrichment analysis of the six RBPs identified, resulted in 14 significantly enriched GOs (FDR ≤ 5%), of which 8 (57%) are for processes involved in splicing (Figure 5B and Supplementary Table S3).

**Figure 5.**
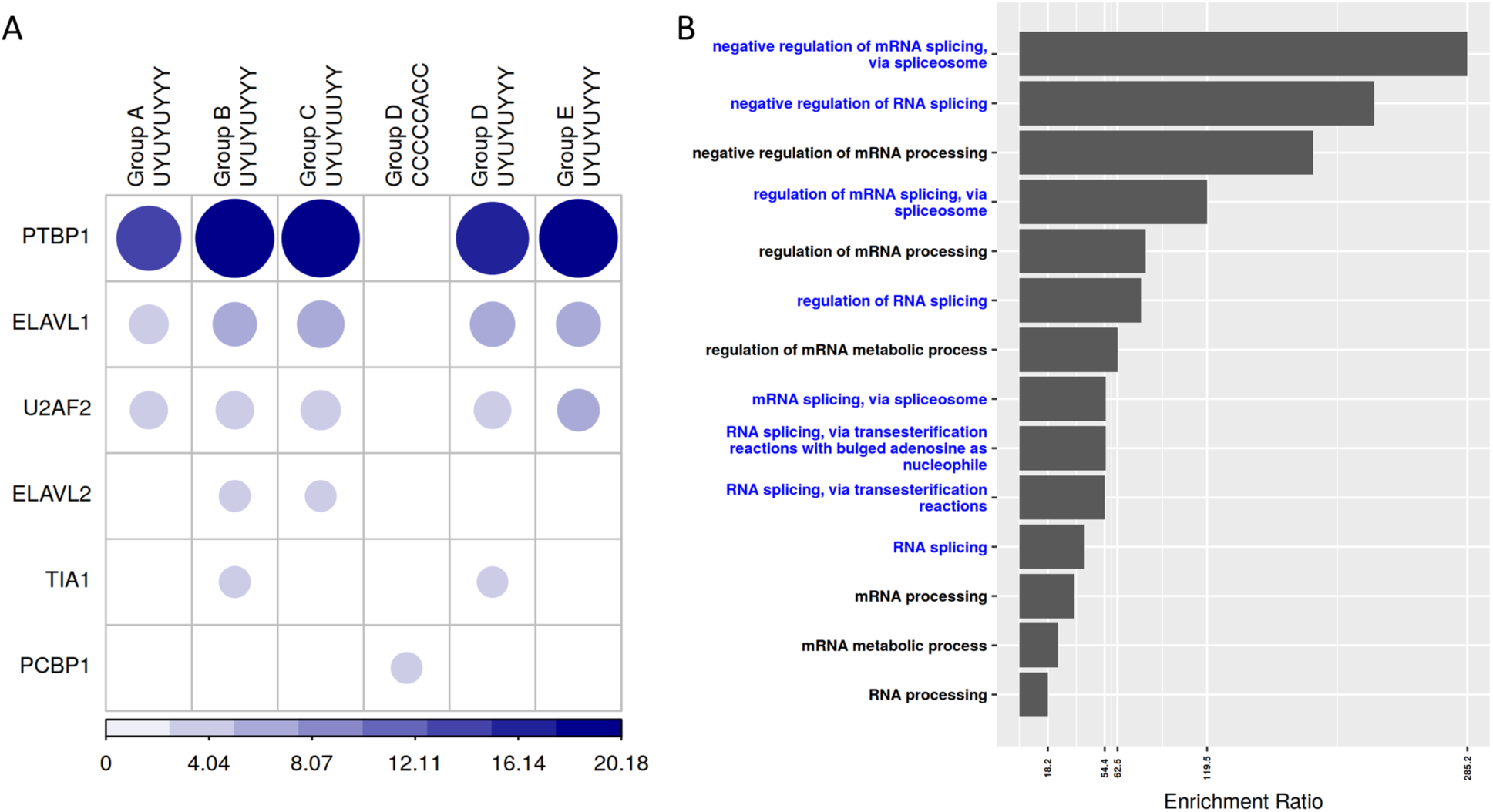
Diagram to represent the similarities of each predictive-base-containing enriched motif with the respective RBP RNA-binding-motifs. **(A)** Presence of the circle indicates that, for that group, the similarity between the enriched motif containing a predictive base identified by our CNN model (indicated at the top) and the known RNA-binding-motif of the RBP indicated at left has reached the adjusted E-value threshold < 0.05. The circle colors and sizes are proportional to the degree of significance (−log E-value) of the sequence similarity, with the values indicated in the scale at the bottom. **(B)** GO biological processes (n = 14) significantly enriched (FDR ≤ 5%) among the six RBPs identified as involved in micro-exon splicing. The x-axis bar represents the enrichment rate (observed/expected) and the y-axis shows the GO categories. The names in blue are for the GOs related to the splicing process.

**Table 3.**
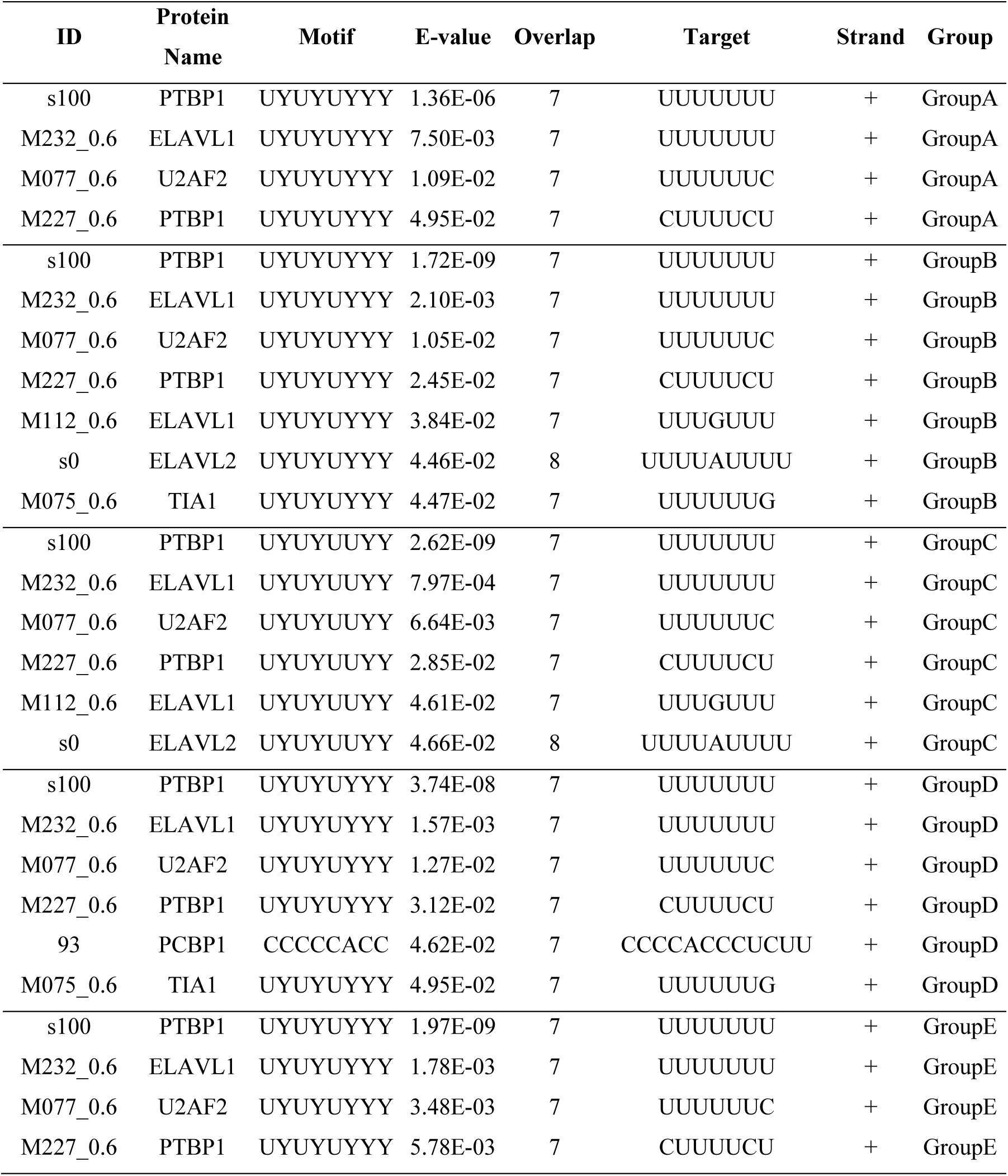
Similarity between the enriched motifs in the introns containing predictive bases and the RBP RNA-binding-motifs

### 3.6 Introns that flank long exons had a different pattern of splicing predictive bases distribution and different conserved RNA-binding-motifs

For comparison, similar analyses were performed with *PositionScores* of bases in the introns flanking long exons (> 39 nt). A heatmap of *PositionScore* values along the introns that flank all tested long exons was generated (Supplementary Figure 3), with the long exons being clustered according to the pattern of *PositionScores* across their flanking introns. Analysis of the more abundant motifs in the intron sequences showed that G and C content or A-rich regions were present (Supplementary Figure 4 and Supplementary Table S4), however the number of sequences comprising each of these motifs was lower than 5 % of total (Figure S4A and Supplementary Table S4), showing that a different pattern of *PositionScore* distribution was found for predictive bases in the introns that flank long exons compared with the pattern in the introns that flank micro-exons.

When these motifs were compared with the ATtRACT database, we found that the motifs in the introns flanking the long exons resulted in the identification of fifteen RBPs (Figure S4B), and all motifs excepted for PCBP1 were different than those identified in the introns flanking the micro-exons.

### 3.7 Intronic splicing silencer motif was enriched in the introns that flank micro-exons

Other databases interrogating conserved RNA sequences were used to explore whether enriched sequences containing predictive bases could harbor additional regulatory region patterns. For this, two other databases were added to the analysis, one for the ISE motifs and one for the ISS. In the ISE database, none of the consensus sequences found in the introns that flank micro-exons reached the E-value similarity threshold ≤ 0.05. It is very interesting to note that in the ISS database, the AGUAGG consensus sequence showed similarity with GroupA Motif 3 (GGRGGAGG, E-value = 0.0175). This motif had not been identified with statistically significant similarity to any RBP motif, in our previous analysis with the ATtRACT database.

### 3.8 Motifs containing predictive bases showed occupancy distribution along flanking intronic regions similar to the distribution of RBP-binding-motifs and ISSs

In order to investigate whether the RBP motifs identified in the previous analysis were represented at a specific location in the upstream or downstream (100 nt) intronic regions that flank micro-exons, the CentriMo tool was used (Bailey and MacHanick, 2012). To perform this analysis, the sequences were divided into upstream and downstream of the exonic/micro-exonic region, and the significant enrichment (E-value ≤ 0.05; Fisher exact-test < 0.05) of each RBP motif in the introns that flank micro-exon compared with the same motifs in the long exon model was calculated and plotted (Figure 6A). In GroupA, four RBP RNA-binding-motifs were identified as showing significant enrichment in the upstream region, namely two different PTBP1 motifs, ELAVL1 and U2AF2 (Figure. 6A), while only PTBP1 motif was enriched in the downstream region (Figure. 6A).

**Figure 6.**
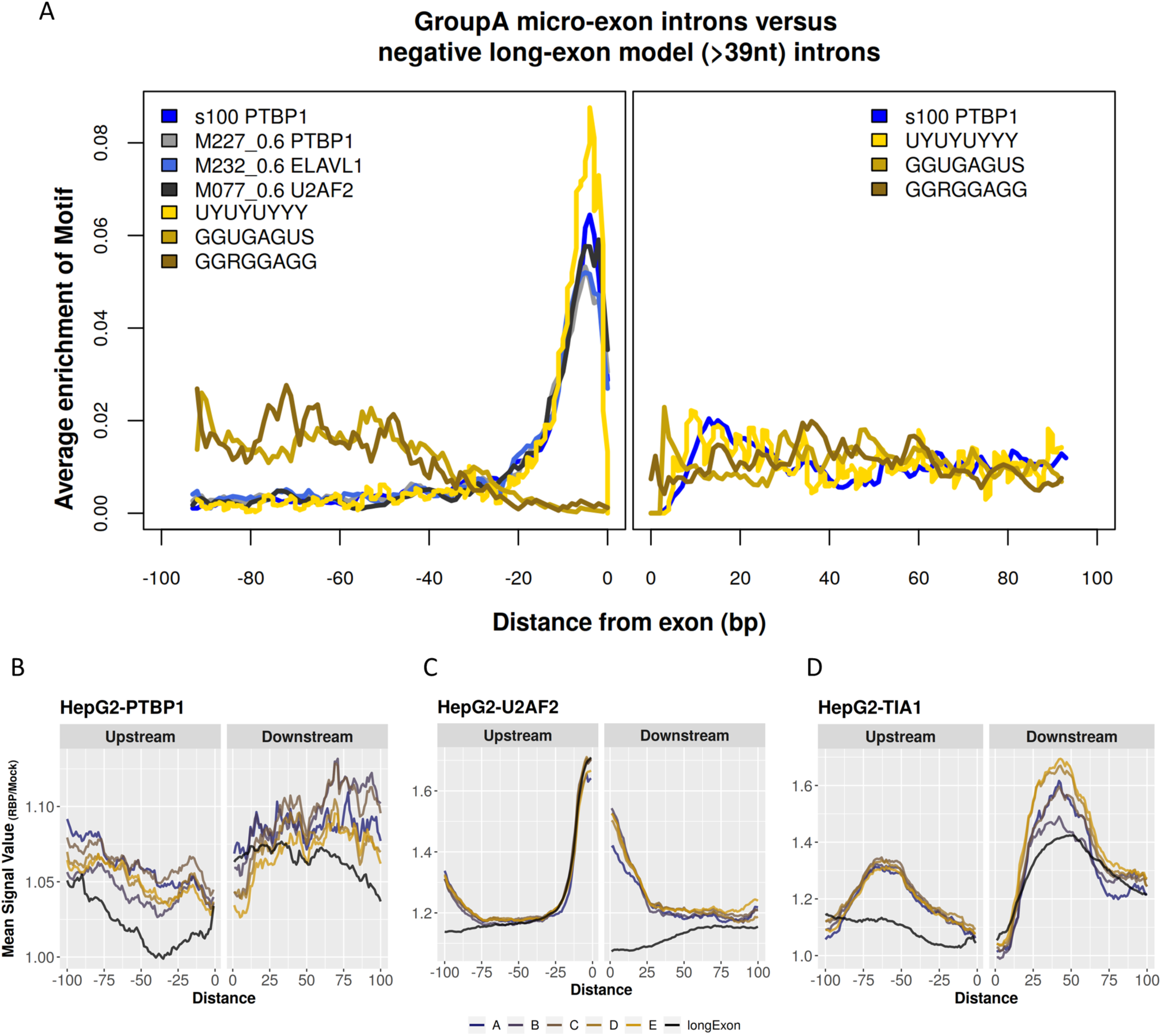
RBP motif enrichment analyses along the intron sequences that flank micro-exons. **(A)** The y-axis represents the average occurrence of the motif along the intronic sequences upstream (−100 to 0) and downstream (0 to +100) of the micro-exons. Data originated from our analysis of enriched motifs that contain micro-exon-predictive bases in the introns that flank micro-exons are gold-colored. All enriched RBPs data from the *in silico* search of RNA-binding-motifs against the ATtRACT Database are plotted with the blue-black color scale. **(B-D)** Re-analysis of eCLIP-seq public data using HepG2 for the analysis of PTBP1 RNA-binding **(B)**, U2AF2 RNA-binding **(C)** and TIA1 RNA-binding **(D)**. The y-axis in B to D represents the average occurrence of signal density for the RBP relative to mock. Signal values are shown with the yellow-brown scale for intronic regions that flank micro-exons of the five groups (GroupA to GoupE), and with the black color for intronic regions that flank long exons (> 39 nt). The x-axis shows the distances along the intron sequence upstream (−100 to 0) or downstream (0 to 100 nt) of the micro-exon (or long exon).

As expected, degenerate Motif 1 UYUYUYYY, which encompasses PTBP1, ELAVL1 and U2AF2 motifs, had a distribution along the intronic sequences neighboring the micro-exons (Figure 6A, yellow line) which was similar to the RBP motifs it represents (Figure 6A, blue lines), while degenerate Motifs 2 and 3 (GGUGAGUS and GGRGGAGG) (Figure 6A, brown lines), which house G-rich regions, showed a completely distinct distribution along intronic upstream regions that flank the micro-exons, with enrichment in the region −40 to −100 nt, away from the micro-exon. Similar patterns of RBP enriched motifs distribution were obtained for all other intronic regions flanking micro-exons in GroupB to GroupE (Supplementary Figure 5).

Next, we performed the distribution analysis of ISS binding motifs along the intronic regions that flank micro-exons. In GroupA, Motif 3 GGRGGAGG harboring a high G content, showed similarity with ISS consensus # I (AGUAGG) and the distribution is shown in Supplementary Figure 6. The distribution suggests that enrichment in the region −40 to −100 nt, away from the micro-exon, was a site for splicing silencers.

### 3.9 eCLIP-seq assays evidenced that PTBP1, U2AF2 and TIA1 bind more abundantly to RNA introns that flank micro-exons compared with long exons

To confirm the above *in silico* findings with experimental approaches, we analyzed publicly available experimental eCLIP-seq data for PTPB1, U2AF2 and TIA1, obtained with two cell lines, namely HepG2 liver carcinoma and K562 leukemia cell lines (Van Nostrand et al., 2016). The density of reads in the intronic RNA regions that flank the micro-exons or long exons was calculated by the ratio between the signal abundance obtained with RBP-specific antibody and the signal in the negative control (mock). In HepG2 liver cells, all three RBPs were found to bind more intensely in the intronic RNA regions that flank micro-exons of all five groups (GroupA to GroupE) in relation to those flanking long exons, as shown in Figure 6B to 6D. Interestingly, PTBP1 (Polypyrimidine Tract Binding Protein 1) showed a higher abundance near the 3’ splice site (3’ss) end, in the RNA introns that flank micro-exons compared with long exons, both upstream (−5 to −50 nt) and downstream of the micro-exons (+60 to +100 nt) (Figure 6B), while U2AF2 showed higher binding in the RNA introns flanking micro-exons compared with long exons in the region near the 5’ss end, in the introns upstream (−100 to −75 nt) and downstream of the micro-exons (+1 to +50 nt) (Figure 6C). Lastly, TIA1 was bound more abundantly to RNA introns flanking micro-exons both upstream (−50 to −25 nt) and downstream (+25 to +60) of the micro-exon (Figure 6D). Similar patterns were observed in K562 leukemia cells (Supplementary Figure 7).

### 3.10 RBP knock down evidenced that PTBP1 and U2AF2 predominantly affected micro-exons splicing

We then looked for possible changes in the splicing patterns of micro-exons that might result from *PTBP1* or *U2AF2* gene knock down. For this, we have re-analyzed RNA-seq gene expression data from both K562 and HepG2 cell lines under *PTBP1* or *U2AF2* knock down (Nostrand et al., 2018), using the vast-tools that is sensitive to alternative splicing events, as described in the Methods.

First, the RNA-seq reads of the *PTBP1* silencing assay and its respective control experiment were mapped to the known splicing junctions in the human genome, and the splicing events detected in both datasets were quantified. A total of 145,836 and 155,528 splicing events were identified in K562 and HepG2, respectively, including intronic retention, alternate use of exons and alternate use of the 3’ and 5’ splice sites. Then, we calculated the *Percent Spliced-In* (PSI) ratio of isoform abundances at each junction and kept those with at least PSI = 0.15 between isoforms. The statistical significance of the difference between the two datasets was calculated using the vast-tools package approach. A total of 208 splicing events in K562, and 258 in HepG2 were identified with significantly altered abundance between the samples when *PTBP1* splicing factor was silenced, compared with the controls. Both analyses showed an enrichment of micro-exon modulation upon *PTBP1* knock-down. Of 208 splicing events in K562 cells, 20 were micro-exon splicing (8.6%, Fisher’s test p-value 1.08E-03, [OR] = 2.28) and of 258 splicing events in HepG2, 35 were micro-exon splicing events (12.15 %, Fisher’s test p-value 1.65E-08, [OR] = 3.26). Most of the changes were related to an increased percentage of micro-exon retention, namely 17 (out of 20, or 85 %) in K562 and 28 (out of 35, or 80 %) in HepG2 when *PTBP1* was silenced (Supplementary Table S5, Supplementary Figure 8A).

Next, in the *U2AF2* silencing assay, a total of 94,568 and 151,369 splicing events were identified in K562 and HepG2 cells, respectively. A total of 977 splicing events in K562, and 1,005 in HepG2 cells were identified with altered abundance between the samples when U2AF2 splicing factor was silenced, compared with the controls. Of these splicing events, 39 were micro-exon splicing events (4 %, Fisher’s test p-value 3.44 E-01, [OR] = 1.08) in K562 cells, and 69 (6 %, Fisher’s test p-value 8.77E-04, [OR] = 1.52) in HepG2 cells. Therefore, only in HepG2 cells the silencing of *U2F2A* showed an enrichment of micro-exon modulation, however in both cell lineage assays the knock down of *U2AF2* led to an increased exclusion of micro-exons. Thus, 32 micro-exons (out of 39, or 82 %) were excluded in K562, and 45 (out of 69, or 65 %) were excluded in HepG2 when *U2AF2* was silenced (Supplementary Figure 8B and Supplementary Table S6).

Noteworthy, *TIA1* knock down resulted in modulation of few micro-exons. There were 132,389 and 148,328 splicing events screened in K562 and HepG2, respectively. Of these, 179 splicing events in K562, and 85 in HepG2 were identified with altered abundance between the samples, compared with control. Only one (0.4 %, Fisher’s test p-value 9.82 E-01, [OR] = 0.25) and 6 (6.4 %, Fisher’s test p-value 1.75 E-01, [OR] = 1.64) splicing events were related to micro-exon modulation in K562 and HepG2, respectively, when *TIA1* was silenced. Thus, for these two cell lines, the knock down of *TIA1* showed little alteration in micro-exon isoforms.

### 3.11 Silencing *PTBP1* modulates the splicing pattern of dystrophin (*DMD*) gene at micro-exon 78 (32-nt-long) on chrX:31,126,642-31,126,673

Since knock down of *PTBP1* showed a predominant modulation of micro-exons in both K562 and HepG2 cell lines, we chose to focus on micro-exon splicing affected by this protein. There were 6 micro-exons that were modulated in common in both cell lines (Supplementary Figure 8A), and one of these was micro-exon 78 of the *DMD* gene (chrX:31,126,642-31,126,673), a 32-nt-long micro-exon. *DMD* is the very long gene that encodes dystrophin, in which deletions of one or many exons cause Duchenne Muscular Dystrophy (OMIM: #310200). The *PositionScores* in the *DMD* intron with the highest negative impact for the micro-exon model were in the upstream region around – 16 nt to – 22 nt (Figure 7A). In fact, this corresponds to one of the intronic regions where PTBP1 binding was found to be enriched in the eCLIP-seq assay (Figure 6B). Knock down of *PTBP1* resulted in an increase in the isoform that harbors this micro-exon in both HepG2 cells (Figure 7B) and K562 cells (Supplementary Figure 9). The likelihood of mean differences in PSI when comparing silencing and control, computed as described by Irimia *et al*. (Irimia et al., 2014) and Tapial *et al*. (Tapial et al., 2017), was 0.13 (at 95 % confidence) in HepG2 cells (Figure 7B, right panel), and 0.12 in K562 cells (Supplementary Figure 9, right panel). The higher abundance of reads mapping to this micro-exon when *PTBP1* was silenced can be clearly observed in Figure 7C, which shows the RNA-seq reads mapping to this genomic locus in each knock down or control assay, for the two cell lines.

**Figure 7.**
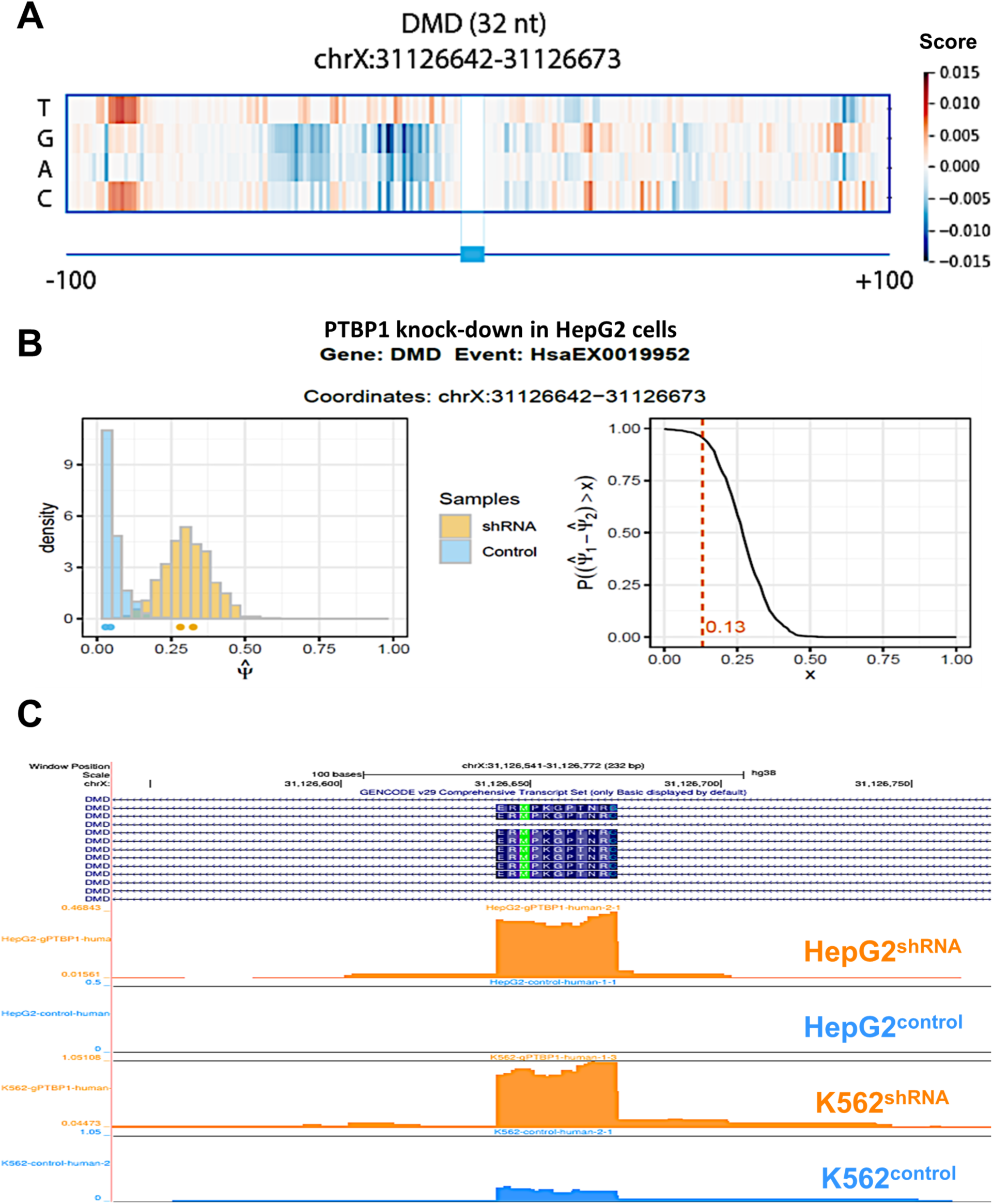
Splicing pattern of dystrophin (*DMD*) micro-exon 78 (32-nt-long) at chrX:31,126,642-31,126,673. **(A)** Positions of critical bases conservation along the introns that flank micro-exon (exon78). The x-axis shows the 200 nucleotides that flank *DMD* gene micro-exon 78 (100 nt at the 5’ or 3’ ends); for each base of the intron sequence that flanks the micro-exon, the delta value (*PositionScore*) of the prediction perturbation caused by the in silico point mutation of that base is represented by the color; the *PositionScore* was calculated by subtracting the intronic sequence prediction value obtained after the base at a given position was in silico mutated to each of the other 3 possible bases from the intronic sequence prediction value using the original wild-type base. Each row represents a specific base, and the base that comprises the wild-type sequence has *PositionScore* = 0 by definition. *PositionScore* color scale is shown at right. **(B)** *DMD* gene micro-exon (32-nt-long) fractional abundance change upon knock down of the *PTBP1* gene in HepG2 cells. The graph on the left shows the density distribution of Percent Spliced-In (PSI) events for the *DMD* gene exon 78. There were only two experimental samples in each of the control (blue) and *PTBP1*-silenced (orange) groups; the curves represent the density distribution of values that the group mean PSI can assume, corrected by the variance of other events that have close PSI values. The graph on the right shows the calculation of the difference between the PSI mean of each of the groups. In this case, there is a 95% probability that the mean difference is 0.13 (ΔPSI = 0.13) between the groups, which means that *DMD* gene micro-exon retention had increased by 13 % upon silencing of the *PTBP1* splicing inhibitor, compared with control. **(C)** Genome browser representation of exon 78 (32-nt-long) locus on Chr. X plus 100 nt intronic sequence on both sides of the exon, in the hg38 assembly. All isoforms of GENCODE annotation for the *DMD* gene are represented in dark blue lines, micro-exon encoded amino acids are represented by dark blue squares. The RNA-seq reads from the *PTBP1* silencing assays that mapped to the locus, from both HepG2 and K562 cell lineages are marked in orange for *PTBP1* knock down shRNA samples, and in light blue for control samples. Only one replicate sample that showed the highest expression of *DMD* exon 78 for each cell line and condition were represented in the Figure.

### 3.12 Multiplexed functional assay of splicing using minigene reporter confirmed that the CNN model can discriminate bases important for micro-exon splicing events

In order to highlight our micro-exon prediction CNN model as a tool to point out nucleotide bases that could affect micro-exon splicing events, we searched the dataset of splice-disrupting variants (SDVs) provided by the work of Cheung *et al*. (Cheung et al., 2019), which employed the Multiplexed Functional Assay of Splicing using Sort-seq (MFASS). In this study, Cheung *et al*. used MFASS to detect splicing event disruption caused by rare genetic variants (Cheung et al., 2019), and screened for 27,733 exonic and intronic single-nucleotide rare variants identified in the Exome Aggregation Consortium (ExAC) database. The authors constructed a synthetic oligonucleotides library that encodes each candidate exon and surrounding intronic sequences with the rare variants (intronic or exonic), and measured the splicing inclusion/exclusion by cloning the synthetic library inside a splicing reporter minigene housing the GFP and mCherry, plus the synthetic sequence flanked by *DHFR* or *SMN1* intron backbone, and integrated the constructs into HEK293T cells using site-specific single-copy integration. If the synthetic exon were excluded, causing an exon skipping, GFP was expressed, otherwise if the synthetic exon were included the mCherry was expressed (Cheung et al., 2019). In total, the work identified 1,050 variants (out of 27,733, i.e. 3.8 %) which were classified as splice-disrupting variants (SDVs) that led to almost complete loss of exon recognition (Cheung et al., 2019), and 6,469 variants (23 %) that caused alteration of ΔPSI ≥ 0.1.

Of the 27,733 variants assayed by Cheung *et al*. (Cheung et al., 2019), only 436 were located at intronic regions of micro-exons, and from these, a total of 27 (6.2 %) were classified as SDVs and 133 (30.5 %) caused a ΔPSI ≥ 0.1 comparing mutant and wild-type. From these 436 assayed variants, in the introns that flank micro-exons, we found that 13 correspond to bases that were present in our list of top 5 % most negative *PositionScore* predictive bases, which would most negatively impact splicing of the flanking micro-exons. Out of these 13 variants assayed, 2 (15.4 %) were classified as SDVs, and 6 (46 %) had an alteration of ΔPSI ≥ 0.1 comparing mutant and wild-type (Supplementary Figure 10). Extending this analysis to the top 25 % predictive bases detected by our CNN model, there were 72 bases screened, of which 6 (8.3 %) were classified as SDVs, and 24 (33 %) presented alteration in ΔPSI ≥ 0.1 comparing mutant and wild type (Supplementary Figure 10). The rate of confirmation of SDVs among the events predicted by the CNN model (8.3 to 15.4 %) was similar to the overall rate of confirmation of SDVs among all assayed rare variants that flank micro-exons (6.2 %) (Cheung et al., 2019). This result shows empirical evidence that the CNN model pointed to a set of intronic bases important for micro-exon splicing events that were among the set of rare variants that affect micro-exon splicing, as detected by large-scale screening with a minigene reporter assay.

## 4 Discussion

In this work we have built a deep learning model using a CNN architecture to identify conservation patterns of intronic DNA sequences important for the micro-exon splicing mechanism, being the first machine learning approach to identify conservation patterns that discriminate micro-exon splicing from long exons splicing. Deep learning methodologies have been extensively used in the genomic context (Poplin et al., 2018), because the algorithm can work with high dimensionality data (input), using layers of spatial abstract features with the combination of multiple kernels, making it possible to handle highly complex data in a hierarchical way (Angermueller et al., 2016; Jones et al., 2017). The original approach of our strategy was to use intronic-flanking base conservation scores among vertebrates combined with *in silico* point mutations of these bases to estimate the impact of intronic mutations on the neighboring micro-exon-splicing predictive power of a CNN model. A delta score value was computed, which was summarized into a *PositionScore* per base in the intron; the largest the absolute *PositionScore* value the higher its impact on the CNN model predictive power. The micro-exon-splicing prediction accuracy of 0.71 obtained with the trained CNN model, and the area under the ROC curve of 0.76, indicated that the performance obtained here was similar to that of other splicing prediction algorithms, such as SpliceRover (Zuallaert et al., 2018), GeneSplicer (Pertea, 2001) and SpliceAl (Jaganathan et al., 2019), which were focused on predicting donor and acceptor splice sites. Of note, none of these approaches (Pertea, 2001; Zuallaert et al., 2018; Jaganathan et al., 2019) did exclusively look for micro-exon splicing patterns.

Micro-exon-flanking intron bases with the highest interspecies conservation values and the smallest, most negative *PositionScore* values were enriched near the micro-exon ends, and the sequence patterns within these regions did possess similarity to known RNA-binding-motifs of RBPs known to affect the splicing mechanism. For example PTBP1 (Li et al., 2015) presents splicing inhibitory properties (Gonatopoulos-Pournatzis et al., 2018) and its silencing in N2A neuroblastoma cells increased the inclusion of 92 % out of 141 altered micro-exons (Han et al., 2014) and higher inclusion of a 12-nt micro-exon in the *KDM1A* gene (Xue et al., 2013). Our re-analyses of the RNA-seq data on the effect of *PTBP1, U2AF2* and *TIA1* splicing-factor genes knockdown in K562 and HepG2 cells (Nostrand et al., 2018), evidenced that among the three factors the largest effect of knockdown was for *PTBP1*, resulting in inclusion of micro-exons due its negative regulatory function (Gonatopoulos-Pournatzis et al., 2018), while *U2AF2* knockdown resulted in an increase of micro-exon skipping. U2AF2 is well characterized to bind to 3’ss splicing enhancer regions (Graveley et al., 2001), being part of a complex important to bind enhancer of micro-exons (eMIC) regions (Faraway and Ule, 2019). This protein mediates splicing of Alu elements in antisense orientation binding to poly-U tracts (Sibley et al., 2016) and is enriched in regions upstream of alternative micro-exons (Li et al., 2015).

Importantly, other RBPs identified in our *in silico* analyses may appear as enriched in experiments using cell lines other than HepG2 and K562, which were used in the public eCLIP-Seq (Van Nostrand et al., 2016) and RBP splicing-factors knockdown (Nostrand et al., 2018) experiments, considering that our *in silico* CNN approach analyzed all the intronic sequences and their interspecies conservation, irrespective of the expression patterns of different splicing-proteins in different tissues/lineages.

Our deep learning CNN model was able to screen at least four bases with negative *PositionScores* in the region –16 nt to –22 nt in the intron upstream of micro-exon 78 of *DMD* gene, indicating that mutations in these bases decreased the likelihood of the intron being correctly classified as a micro-exon-flanking region. Splicing of micro-exon 78 (32-nt-long) of the *DMD* gene is an example of a micro-exon splicing event in a human gene involved with a debilitating and lethal disease, the Duchenne Muscular Dystrophy, which results from mutations that cause splicing errors (Le Rumeur et al., 2010). The *DMD* gene is the longest gene in the human genome, which spans over 2.2 Mb, with long introns that are processed through non-sequential and multi-splicing steps (Gazzoli et al., 2016), and in this context the correct mechanism of splicing is essential. Indeed, splicing defects in the *DMD* gene have been identified as originated from mutations both at canonical sites and located at less-conserved positions deeply embedded within the large *DMD* introns (Tuffery-Giraud et al., 2017). Different alternatively spliced isoforms of *DMD* are expressed in diverse tissues such as skeletal muscle, brain and smooth muscle (Feener et al., 1989). Also, the Dp71 transcript, encoding a 70-75 kDa C-terminal protein product of the *DMD* gene expressed in the human brain (Austin et al., 2000), shows several isoforms with alternative C-terminal, including one with exon 78 skipping, which changes the reading frame and modifies the translated C-terminal, producing dystrophin with a 31 amino acids (aa) tail instead of a shorter 13 aa tail (Austin et al., 2000). Dysregulation of these splicing isoforms were related to cognitive impairments (Tadayoni et al., 2012), although the mechanisms of dysregulation are not known. Regarding specifically the isoform without exon 78, it is expressed in embryonic stages in pre-contractile muscle, and re-expression of this isoform instead of the adult isoform contributes to progression of the dystrophic process in myotonic dystrophy type I (Rau et al., 2015). On the other hand, the isoform with exon 78 skipping is the most expressed in neuronal SH-SY5Y cells (Nishida et al., 2015), while in muscle tissue under physiological conditions, only 2.5 % of the expressed gene corresponds to this alternative isoform (Tuffery-Giraud et al., 2017). All this suggests that the correct expression of *DMD* isoforms is under developmental control and must involve a complex machinery; we speculate that mutations in the conserved region around bases –16 nt to –22 nt upstream of micro-exon 78 might affect its fine-tuning splicing regulation.

Overall, the deep learning CNN model has pointed to intron bases which had a high predictive value for micro-exon splicing, and a search for conserved patterns has identified RNA-binding-motifs of specific RBPs associated with the splicing process. Even more interesting was the finding that the *in silico* motif predictions could be experimentally confirmed with data from e-CLIP RBP binding assays, from silencing assays of splicing-regulatory proteins, and from splice-disrupting mutations detected with minigene reporter, thus reinforcing the predictive power of the *in silico* model. Search for the impact of variants on splicing mechanism has gained attention during the past years (Li et al., 2016), which resulted from gathering information about sQTL in the human population (Park et al., 2018) or in disease (Tian et al., 2019); especially considering the noncoding regions, it is still a challenge to discriminate risk variants outside of exon regions in complex diseases (Xiao et al., 2017).

RNA-seq deep-sequencing has been frequently used to assemble novel transcripts, and the predictive power of our CNN model could be applied as a tool to validate *de novo* micro-exons annotation in different tissues or cancer cells.

Finally, we propose that the deep learning CNN model developed here could be used in combination with the full genome sequencing data from patients, in order to perform an unbiased screening for mutations in the intronic regions of genes, looking for point mutations in bases that have high impact on micro-exon-splicing predictive power. This approach can possibly reveal critical point mutations in intronic conserved regions that flank micro-exons that would be related in a yet unknown manner to a given disease.

## 6 Author Contributions

LFdS and SVA conceived the work. LFdS, ACT and VM performed the experiments and obtained the data. LFdS, ACT and SVA analyzed and interpreted the data. LFdS and ACT wrote the first draft of the manuscript. SVA edited the draft and wrote the final manuscript. All authors read and approved the final manuscript.

## 8 Acknowledgments

We thank Dr. J.C. Setubal for access to the computational facilities of the Bioinformatics Laboratory of Instituto de Química, Universidade de São Paulo (USP).

## 1 Data Availability Statement

The datasets analyzed during the current study are available in the ENCODE repository, at https://www.encodeproject.org.

## Supporting Information

**Supplementary Figure 1.**
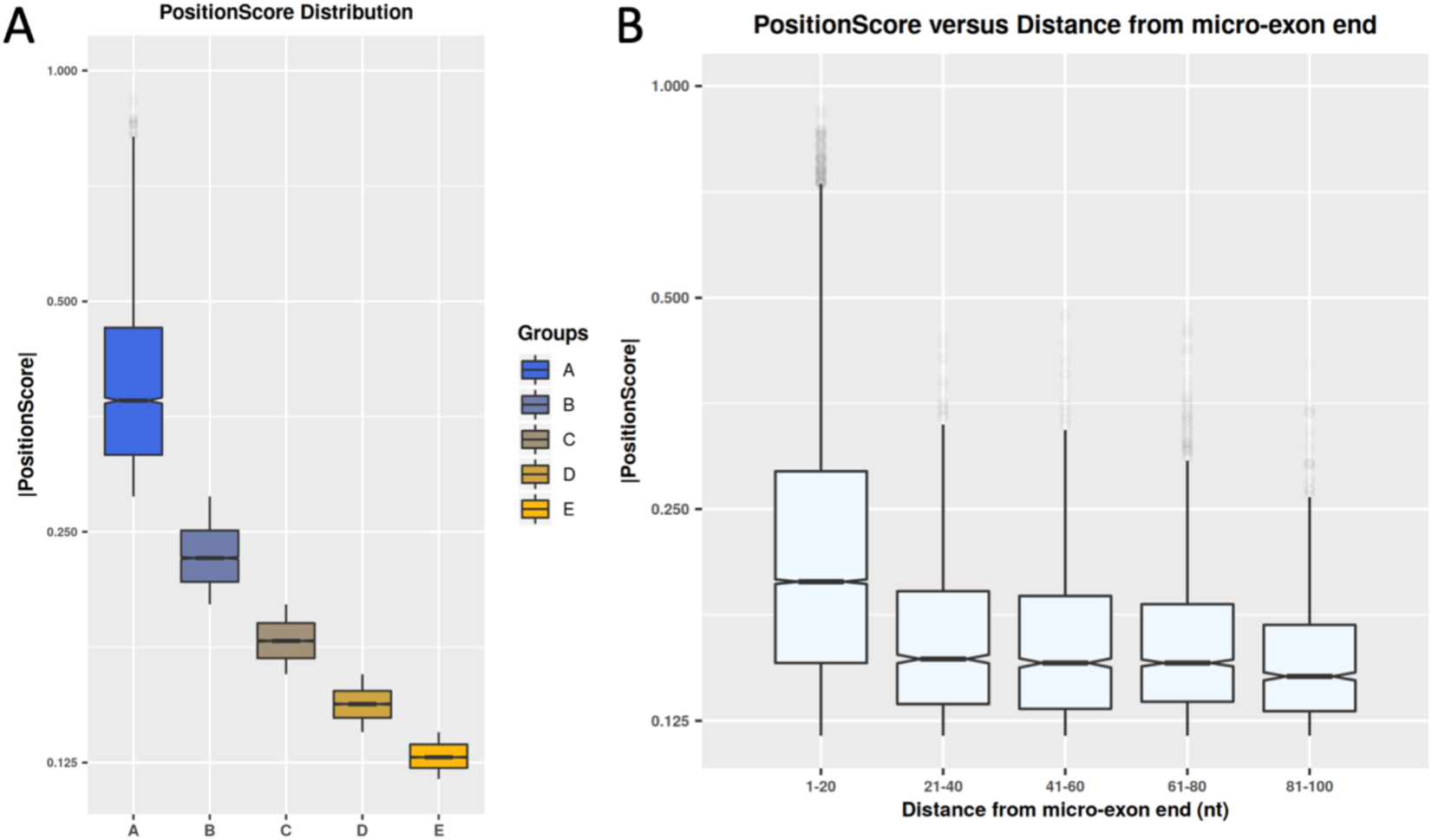
*PositionScore* absolute values across the five groups and their values versus distance to micro-exon end. **(A)** Boxplot distribution of *PositionScore* values within each of the five groups (A to E) into which the top 5 % most predictive intron bases identified by the Deep Learning CNN algorithm were divided. *PositionScore* values were calculated for each of the 200 bases that flank the 5’- and 3’-end of each of 4,908 micro-exons, and the top 5 % most predictive bases (with the highest *PositionScore* absolute values) were retrieved and divided into 5 groups, GroupA representing the top 1 % highest percentile and GroupE the lowest percentile. The y-axis shows the absolute value of *PositionScore*, the x axis shows the five different groups. **(B)** Box plot of absolute values of *PositionScore* of intron predictive bases (y-axis) as function of the distance to the micro-exon end was computed in 20-nt windows along the intron (x-axis). All *PositionScores* from the five groups (GroupA to GroupE) were plotted together. Correlation between *PositionScore* and distance to the micro-exon was calculated (Kendall’s rank correlation, tau = −0.23, p-value < 2.2e-16).

**Supplementary Figure 2.**
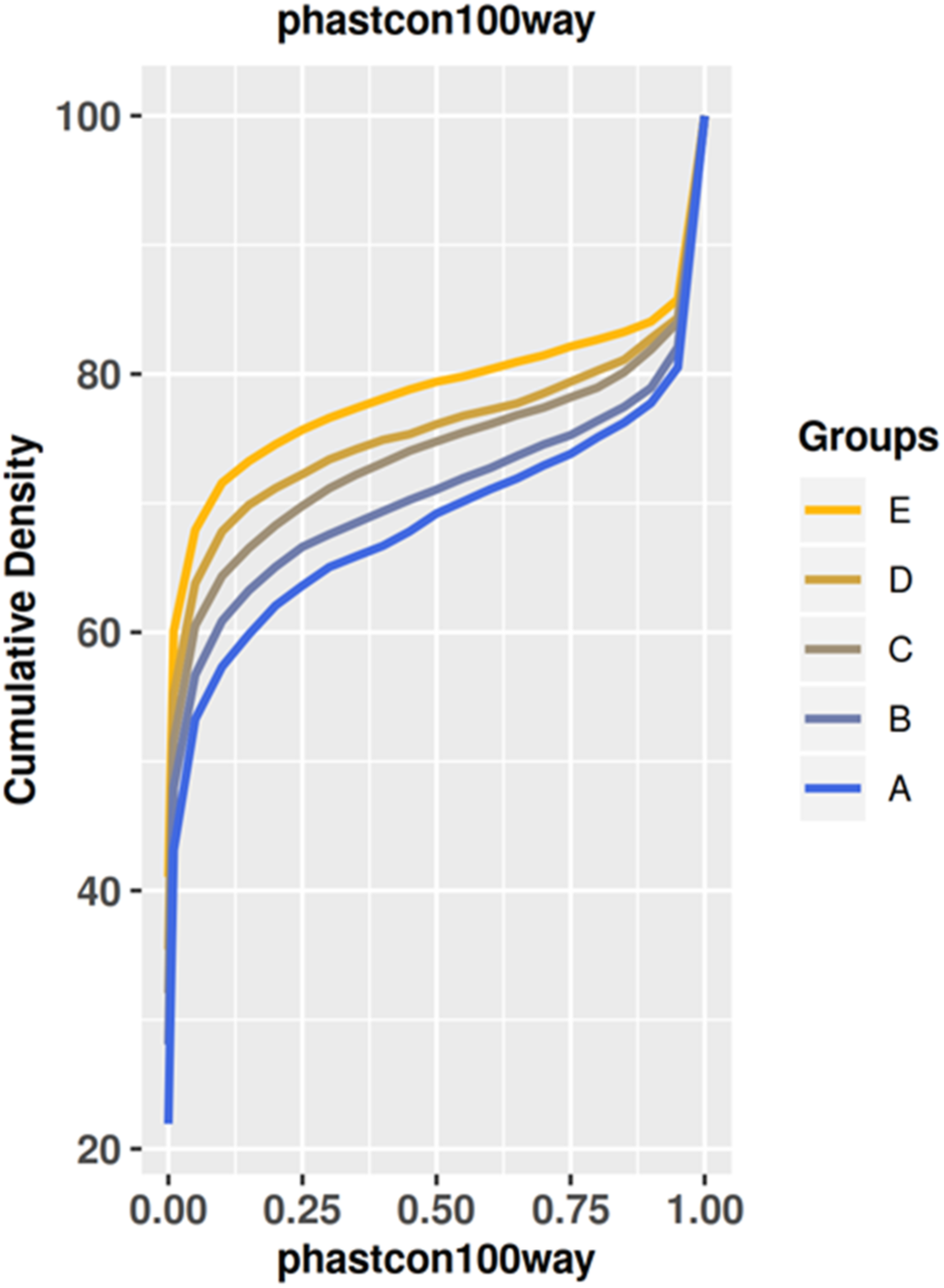
Cumulative Distribution of PhastCon100way according to groups of splicing-predictive bases *PositionScore*. Cumulative distribution of PhastCon100way values for each of the five groups, indicated by colors. The y-axis shows the cumulative distribution and the x-axis shows the PhastCon100way score. Statistical differences in PhastCon100 way scores distribution were observed in all comparisons (KS-test, p-value < 0.05).

**Supplementary Figure 3.**
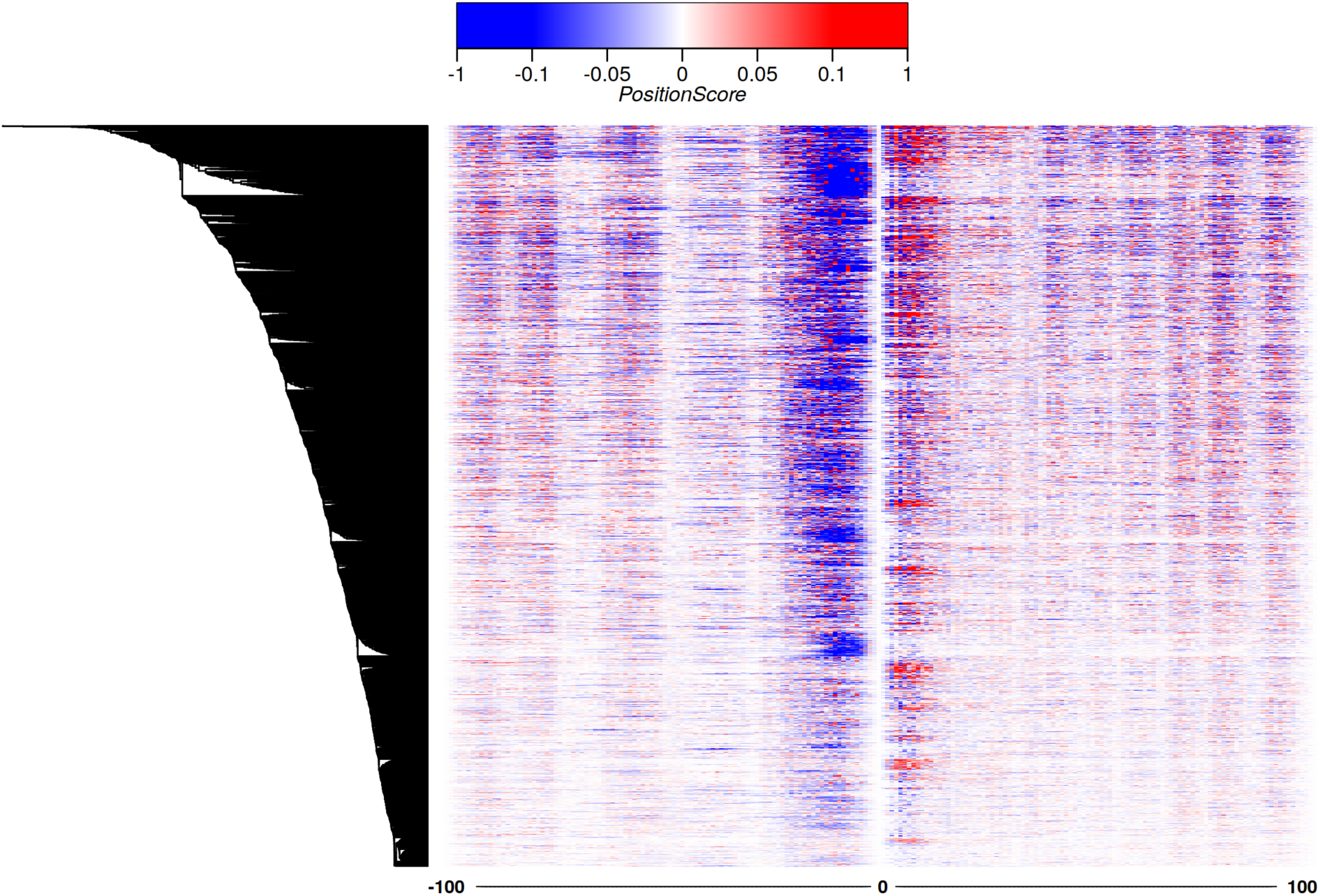
*PositionScores* heatmap of long-exon model introns. On the y-axis each of the 4,417 long exons used in training and validation the Deep Learning CNN model is represented in one line. The x-axis shows the 200 nucleotides that flank each long-exon (100 nt at the 5’ or 3’ ends). The delta value (*PositionScore*) of the prediction perturbation caused by the *in silico* point mutation of that base is represented by the color; the delta is calculated by subtracting the intronic sequence prediction value, obtained after the base at a given position was changed to the other 3 possible bases, from the intronic sequence prediction value using the original wild-type base. The heatmap has clustered the long exons according to the *PositionScore* pattern of the intron sequences that flank each long exon. On the upper left is the color scale of the perturbation *PositionScore* values. Positive values indicate that the *in silico* mutation increased the probability that a given sequence was classified as an intron that flank a long exon, and negative values show that the in silico mutation increased the probability that the sequence was mistakenly classified as an intron that flank a micro-exon.

**Supplementary Figure 4.**
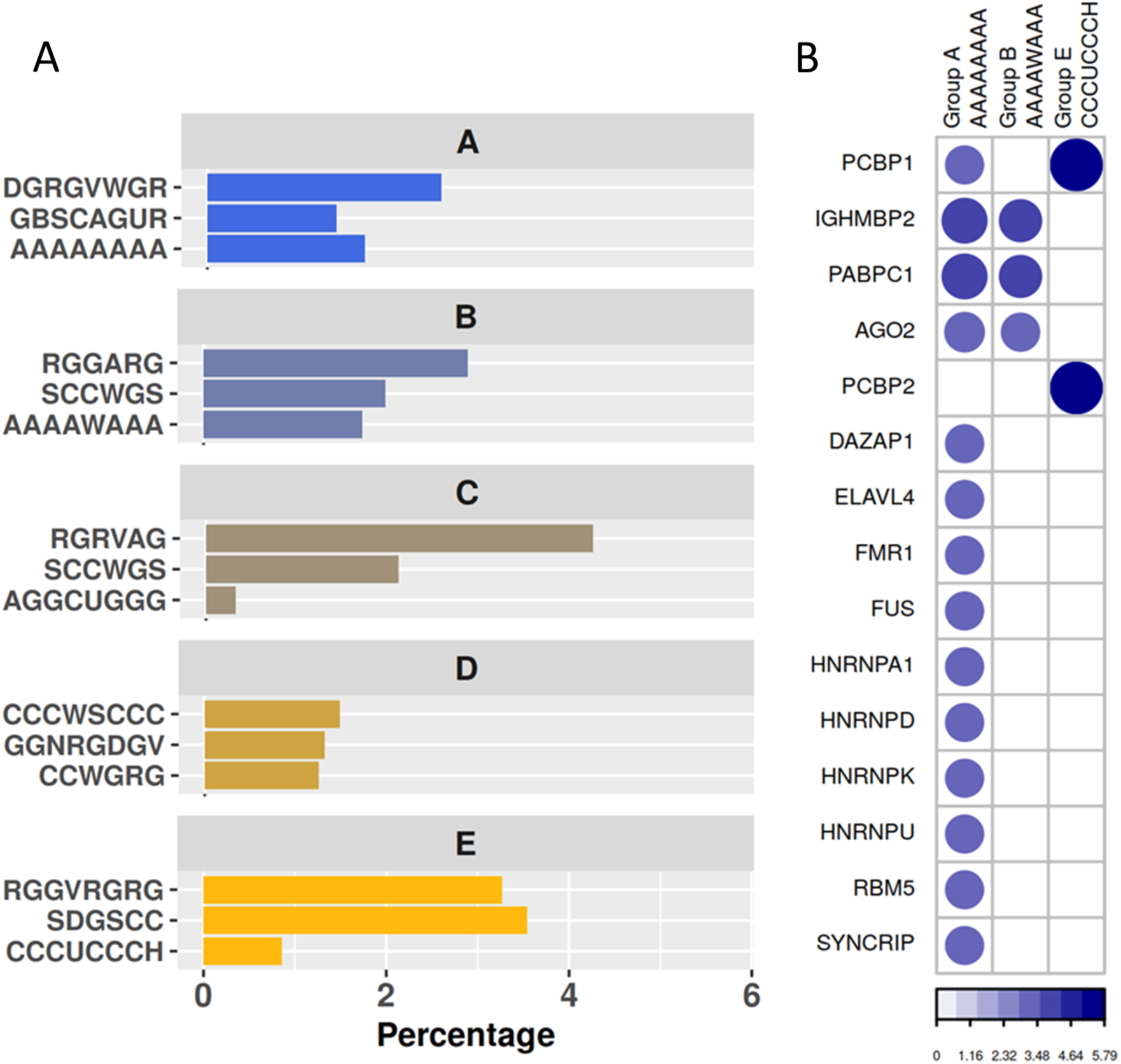
Enriched sequence motifs that contain the predictive bases identified by the CNN model in the introns of neighboring long exons. (A) Percent of intron sequences that support the enrichment of each motif indicated at left. The x-axis shows the percent of intron sequences that contain the predictive bases within the corresponding motif, calculated as a percentage of the total number of sequences in the corresponding group (A to E). The y-axis shows for each group the sequence motifs 1 to 3, ordered from top to bottom according to the enrichment significance (based on the adjusted E-value). (B) Similarities of each predictive-base-containing enriched motif with the respective RBP RNA-binding-motifs. Presence of the circle indicates that, for that group, the similarity between the enriched motif containing a predictive base (indicated at the top) and the known RNA-binding-motif of the RBP indicated at left has reached the adjusted E-value threshold <0.05. The circle colors and sizes are proportional to the degree of significance (−log E-value) of the sequence similarity, with the values indicated in the scale at the bottom.

**Supplementary Figure 5.**
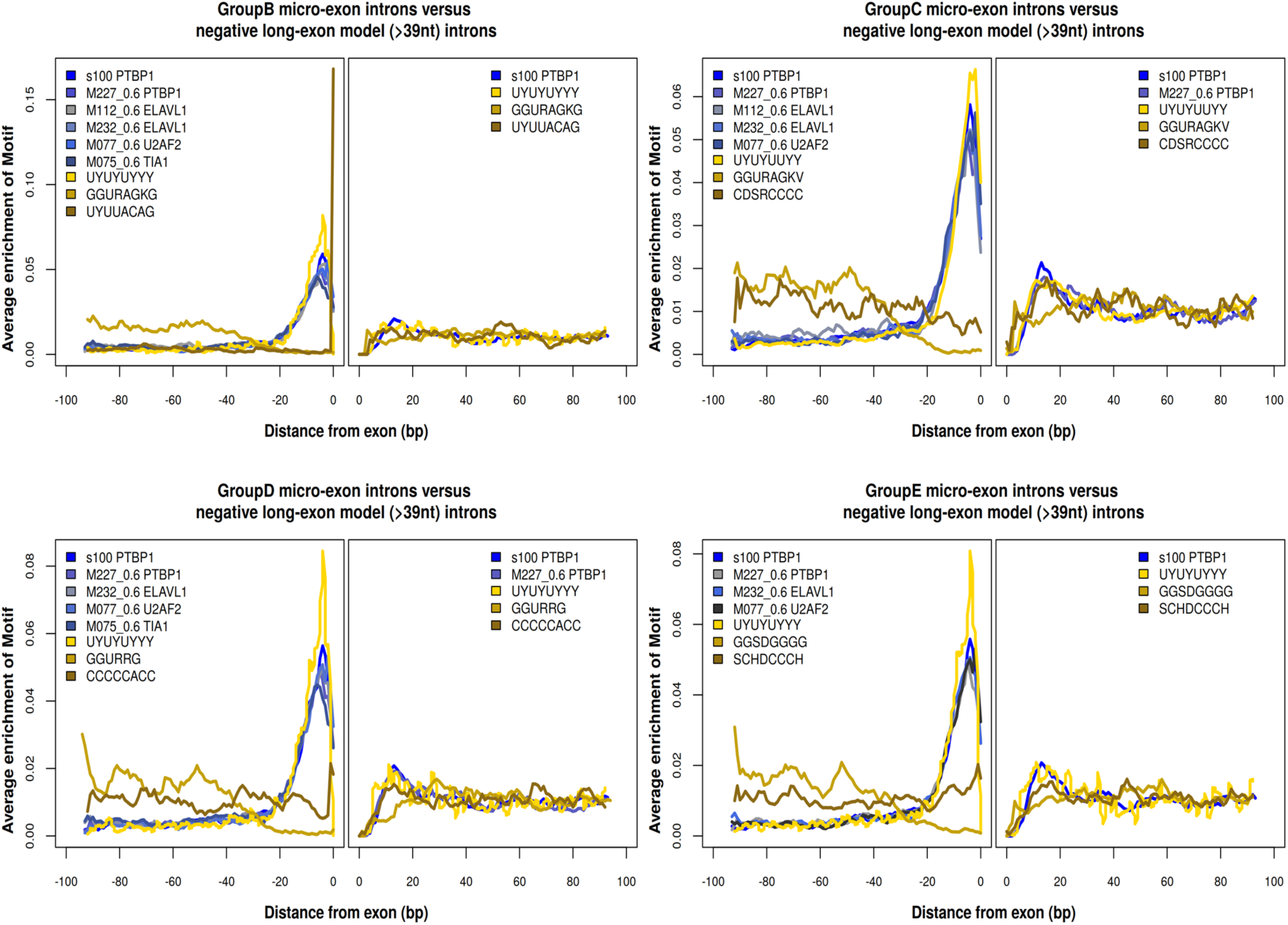
RBP motif enrichment in silico analyses along the intron sequences that flank micro-exons. The y-axis represents the average occurrence of the motif along the intronic sequences upstream (−100 to 0) and downstream (0 to +100) of the micro-exons. Data originated from our analysis of enriched motifs that contain micro-exon-predictive bases in the introns that flank micro-exons are gold-colored. All enriched RBPs data from the *in silico* search of RNA-binding-motifs against the ATtRACT Database (Giudice et al., 2016) are plotted with the blue-black color scale.

**Supplementary Figure 6.**
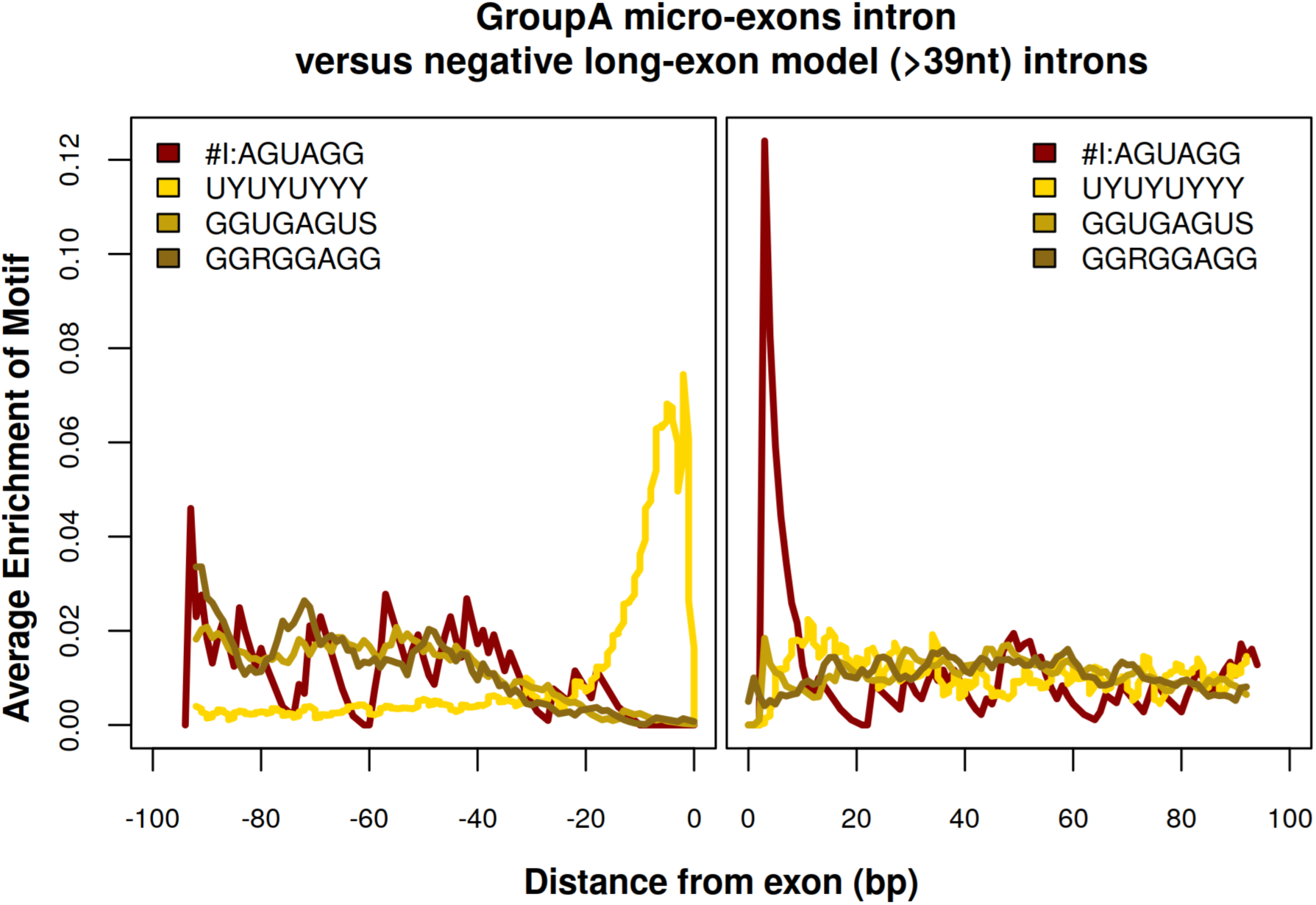
ISS motif enrichment *in silico* analyses along the intron sequences that flank micro-exons. The y-axis represents the average occurrence of the motif along the intronic sequences upstream (−100 to 0) and downstream (0 to +100) of the micro-exons. Data originated from our analysis of enriched motifs that contain micro-exon-predictive bases in the introns that flank micro-exons are gold-colored. All enriched RBPs data from the in silico search of RNA-binding-motifs against the 10 consensus sequences of ISS defined by Wang et al. (Wang et al., 2013b), are plotted with the red color scale; only one sequence showed enrichment, namely ISS_consensus #I (AGUAGG).

**Supplementary Figure 7.**
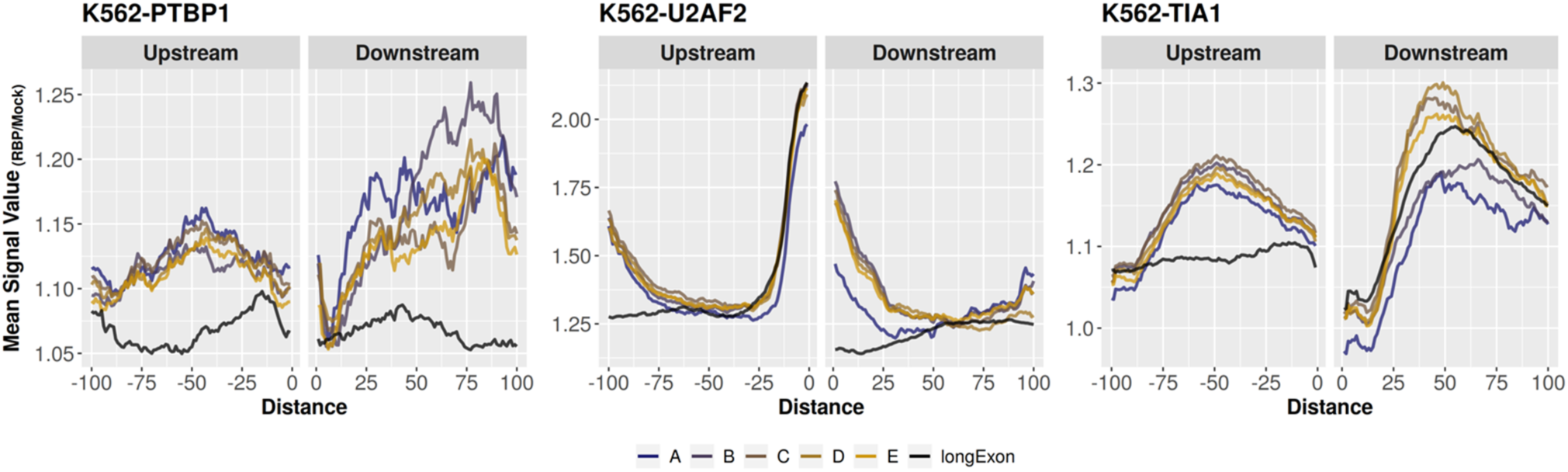
Re-analysis of eCLIP-seq public data using K562 cells for measuring of PTBP1 RNA-binding, U2AF2 RNA-binding and TIA1 RNA-binding. The y-axis represents the mean occurrence of signal density for the RBP relative to mock. Signal values are shown with the yellow-brown scale for intronic regions that flank micro-exons of the five groups (GroupA to GoupE), and with the black color for intronic regions that flank long exons (> 39 nt). The x-axis shows the distances along the intron sequence upstream (−100 nt to 0) or downstream (0 to 100 nt) of the micro-exon (or long exon).

**Supplementary Figure 8.**
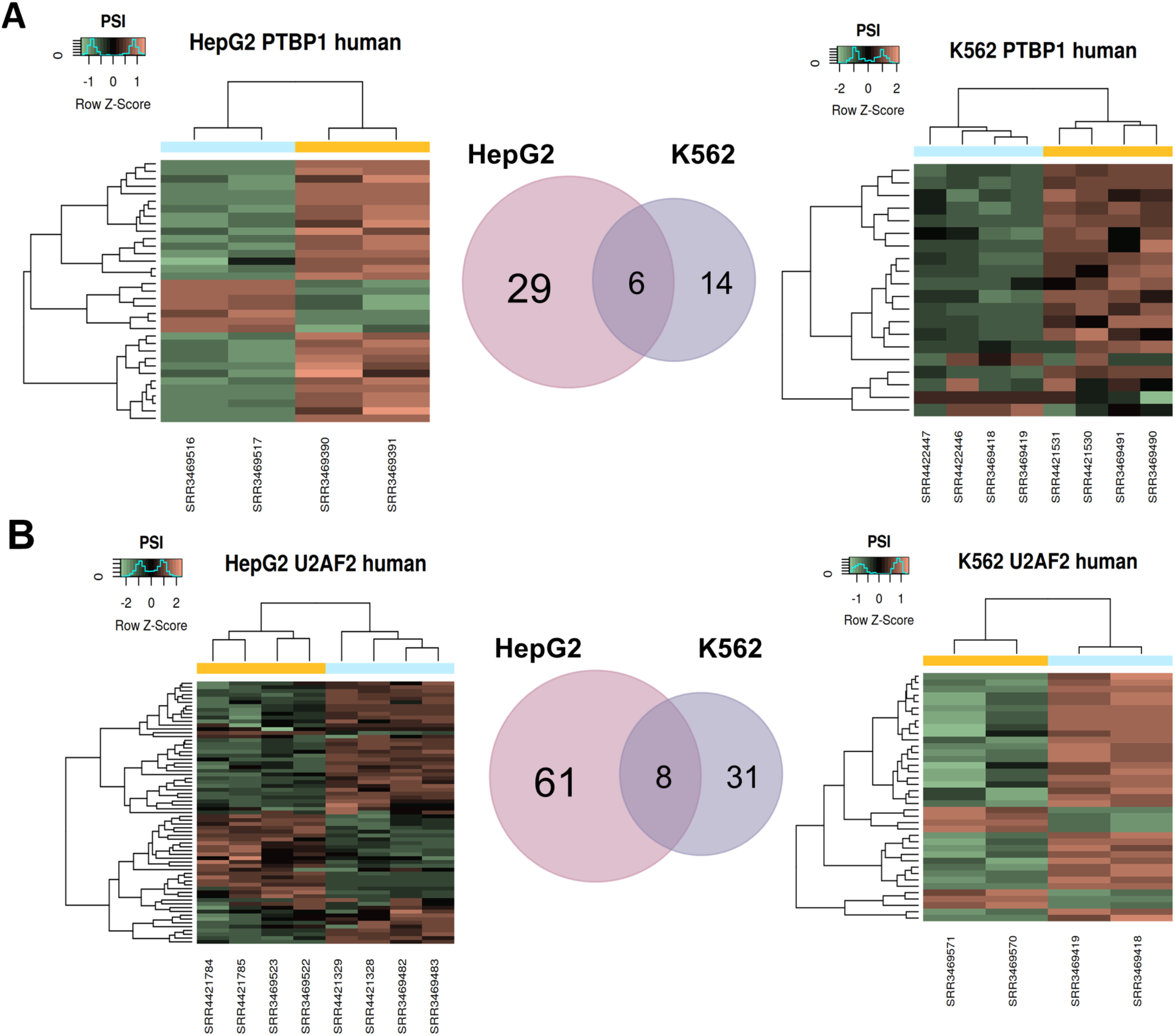
Re-analysis of RNA-seq expression public data for quantification of the effect on micro-exon *percent-spliced-in* (PSI) caused by silencing *PTBP1* (A) or *U2AF2* (B) in HepG2 cells (left) and in K562 cells (right). The y-axis represents micro-exons that were altered when comparing shRNA and control. Signal values are shown with the green-red scale; red represents increase abundance of isoform expression and green low abundance. The x-axis shows the libraries used to perform analysis with vast-tools. All micro-exon events presented a PSI > 0.15 and 95 % probability that the mean difference is |ΔPSI| > 0.10 between the groups. Venn Diagram shows the overlapping micro-exon splicing events in each cell line.

**Supplementary Figure 9.**
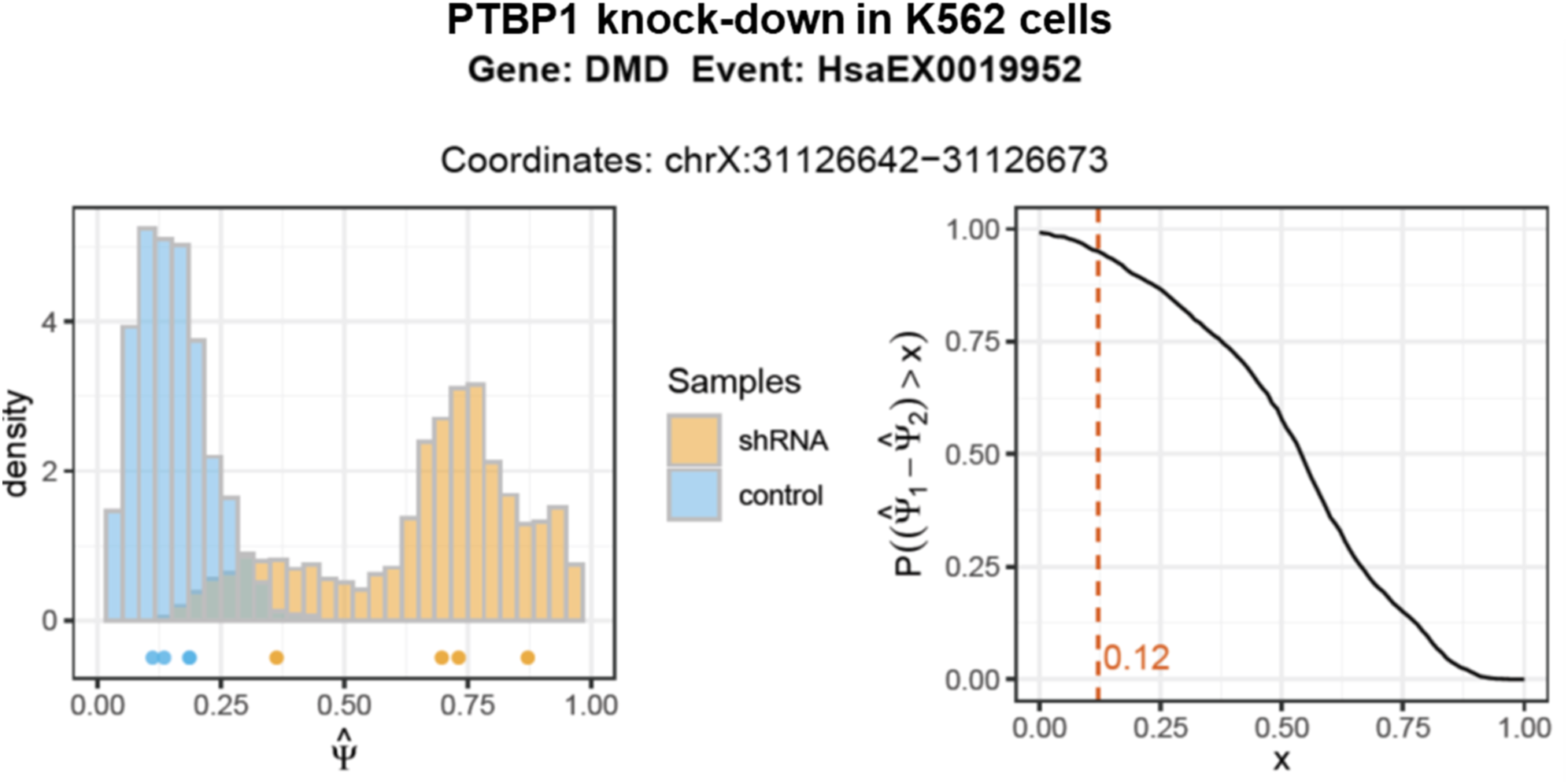
*PTBP1* knock-down in K562 cells caused an increase in *DMD* micro-exon 78 insertion. *DMD* gene micro-exon 78 (32-nt-long) fractional abundance change upon knockdown of the *PTBP1* gene in K562 cells. The graph on the left shows the density distribution of Percent Spliced-In (PSI) events for micro-exon 78 of the *DMD* gene. There were only two experimental samples in each of the control (blue) and *PTBP1*-silenced (orange) groups; the curves represent the density distribution of values that the group mean PSI had assumed, corrected by the variance of other events that had close PSI values. The graph on the right shows the calculation of the difference between the PSI mean of each of the groups. In this case, there was a 95 % probability that the mean difference (ΔPSI) was 0.12 between the groups, which means that *DMD* gene micro-exon 78 retention had increased by 12 % upon silencing of the *PTBP1* splicing inhibitor, compared with control.

**Supplementary Figure 10.**
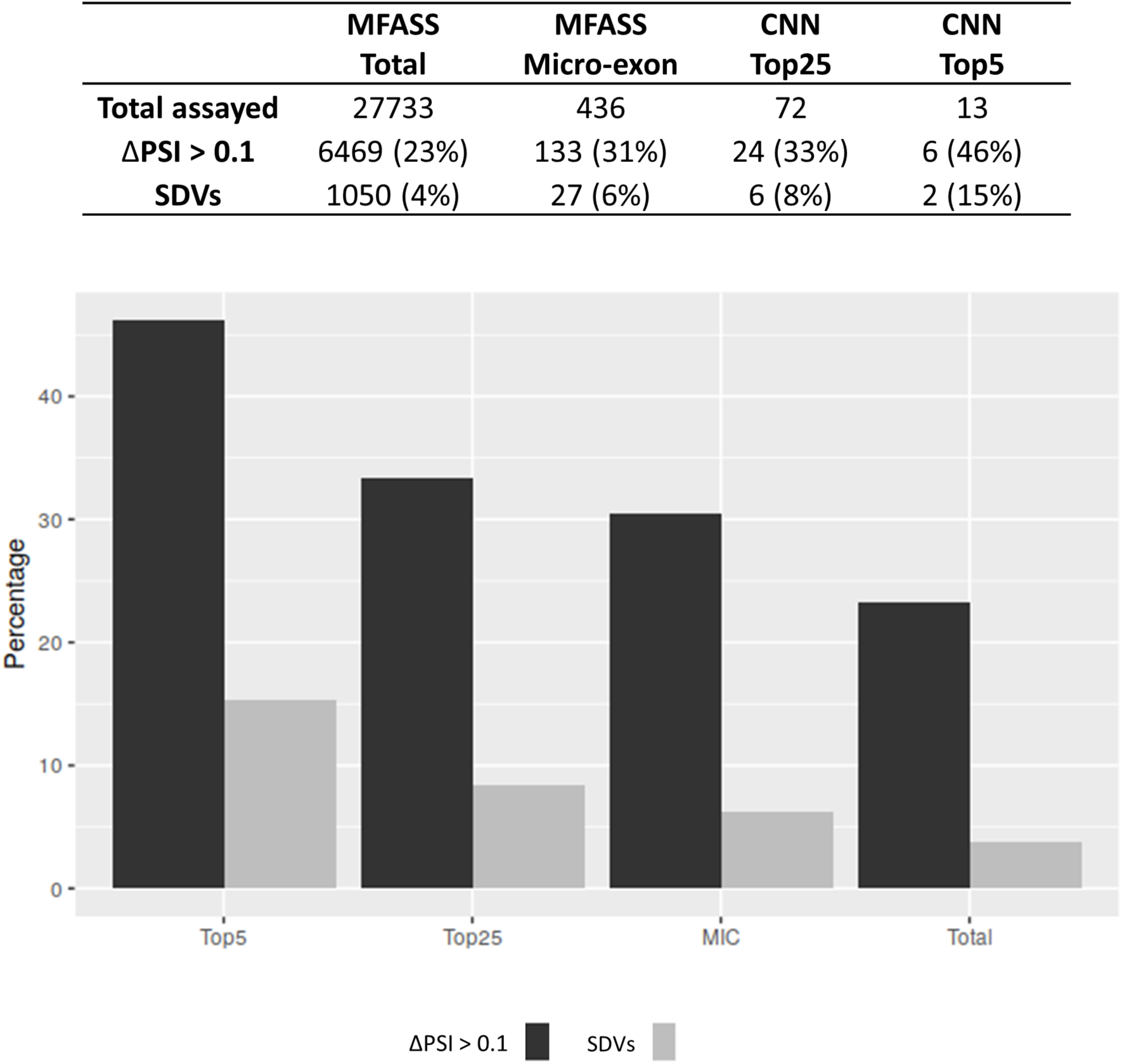
Cross-comparison between the Multiplexed Functional Assay of Splicing using Sort-seq (MFASS) and the percent bases predicted by the CNN *PositionScore* model. The table at the top shows our re-analysis of MFASS data obtained by Cheung et al. (Cheung et al., 2019), for the screening with a minigene reporter of 27,773 rare variants in the human genome that had an effect on splicing. The **first column** shows the total number of variants assayed, the percentage of those that caused a change in splicing isoforms of |ΔPSI| > 0.1, and the percentage that were classified as Splice-Disrupting Variants (SDVs) (|ΔPSI| > 0.5). The **second column** shows the number of variants assayed by Cheung et al. that were in introns that flanked micro-exons. The **third and fourth columns** show the number of variants assayed by Cheung *et al*. (Cheung et al., 2019) in introns that flanked micro-exons, whose bases were also classified by our CNN model as being among the Top25, or among the Top5, with absolute *PositionScores* ranking among the 25 % or among the 5 % with the highest prediction of impacting the micro-exon splicing, respectively. **Bar graph** at the bottom shows the same percentage data shown on the table at the top; MIC are bases screened by Cheung *et al*. (Cheung et al., 2019) that flank only intronic regions of micro-exons; Total represents the group of all bases assayed in the study of Cheung *et al*. (Cheung et al., 2019), including intronic and exonic bases located in micro-exons and long exons.

**Supplementary Table S1. Comparison of distance distribution (nt) using KS-test**

**Supplementary Table S2. Comparison of PhastCon Score using KS-test**

**Supplementary Table S3. Biological Pathways GO enrichment analysis of RBPs**

**Supplementary Table S4. Top three most enriched sequence motifs found in each of the groups of intronic sequences that flank long-exons**

**Supplementary Table S5. Micro-exon splicing events differentially expressed upon *PTBP1* knock down (KD) compared with control**

**Supplementary Table S6. Micro-exon splicing events differentially expressed upon *U2AF2* knock down (KD) compared with control**

